# Genetic Diversity in the IZUMO1-JUNO Protein-Receptor Pair Involved in Human Reproduction

**DOI:** 10.1101/2021.08.08.455586

**Authors:** Jessica Allingham, Wely B. Floriano

## Abstract

Fertilization in mammals begins with the union of egg and sperm, an event that starts a cascade of cellular processes. The molecular-level understanding of these processes can guide the development of new strategies for controlling and/or promoting fertilization, and inform researchers and medical professional on the best choice of interventions. The proteins encoded by the IZUMO1 and JUNO genes form a ligand-receptor protein pair involved in the recognition of sperm and egg. Due to their role in the fertilization process, these proteins are potential targets for the development of novel anti-contraceptive, as well as infertility treatments. Here we present a comprehensive analysis of these gene sequences, with the objective of identifying evolutionary patterns that may support their relevance as targets for preventing or improving fertility among humans. JUNO and IZUMO1 gene sequences were identified within the genomes of over 2,000 humans sequenced in the 1000 Genomes Project. The human sequences were subjected to analyses of nucleotide diversity, selection neutrality, population-based differentiation (F_ST_), haplotype inference, and whole chromosome scanning for signals of positive or of balancing selection. Derived alleles were determined by comparison to archaic hominin and other primate genomes. The potential effect of common non-synonymous variants on protein-protein interaction was also assessed. IZUMO1 displays higher variability among human individuals than JUNO. Genetic differentiation between continental population pairs was within whole-genome estimates for all but the JUNO gene in the African population group with respect to the other 4 population groups (American, East Asian, South Asian, and European). Tajima’s D values demonstrated deviation from neutrality for both genes in comparison to a group of genes identified in the literature as under balancing or positive selection. Tajima’s D for IZUMO1 aligns with values calculated for genes presumed to be under balancing selection, whereas JUNO’s value aligned with genes presumed to be under positive selection. These inferences on selection are both supported by SNP density, nucleotide diversity and haplotype analysis. A JUNO haplotype carrying 3 derived alleles out of 5, one of which is a missense mutation implicated in polyspermy, was found to be significant in a population of African ancestry. Polyspermy has a disadvantageous impact on fertility and its presence in approximately 30% of the population of African ancestry may be associated to a potentially beneficial role of this haplotype. This role has not been established and may be related to a non-reproductive role of JUNO. The high degree of conservation of the JUNO sequence combined with a dominant haplotype across multiple population groups supports JUNO as a potential target for the development of contraceptive treatments. In addition to providing a detailed account of human genetic diversity across these 2 important and related genes, this study also provides a framework for large population-based studies investigating protein-protein interactions at the genome level.

**AUTHOR SUMMARY:** Fertilization in mammals depends on egg and sperm connecting to each other. Two proteins that are essential for this process are egg’s JUNO and sperm’s IZUMO1. These proteins are encoded by genes known as JUNO and IZUMO1, which are present in the genomes of all humans. We analyzed the publicly available genomes of over 2,000 individuals from different continental populations (African (AFR), American (AMR), East Asian (EAS), European (EUR) and South Asian (SAS)) to investigate how well-conserved these genes are among humans, and to look for any signs that genetic variation on these genes could impact the binding of their encoded proteins to one another which, in turn, could impact the fertilization process. We found that the JUNO gene in the African population sampled diverged the most compared to the other population groups, and this divergence was above what is generally observed across the genome. Around 30% of the people of African ancestry investigated here carry a mutation that is associated to the abnormal fertilization of an egg by more than one sperm (known as polyspermy). Why this mutation is so common in the African population should be further investigated. We speculate that this mutation confers a genetic advantage related to the role of this receptor-ligand pair in cells not involved in reproduction.

## INTRODUCTION

Fertilization is a multistep process that includes sperm-egg recognition and sperm-egg fusion, resulting in a genetically distinct diploid organism[1]. Fusion shows less distinct species-specificity than does the sperm-egg recognition step of fertilization, which suggests that the mechanism and molecules responsible for the fusion of the gametes are more highly conserved amongst different species than the ones responsible for sperm-egg recognition[2]. Sperm acquire the ability to fertilize the egg by exposing previously concealed receptor proteins onto their surface following the acrosome reaction[1]. JUNO and IZUMO1 are two proteins whose interaction has been identified as essential in the sperm-egg recognition step of the fertilization process[3]. They are the first essential sperm-egg recognition pair identified in any species, and their ability to bind has been confirmed in many mammals[4].

IZUMO1 is a protein located on the cell surface of capacitated sperm cells, named after a Japanese marriage shrine[5]. It is encoded by a gene located on chromosome 19 in the human genome[6]. IZUMO1 has been shown to play a crucial role in fertilization. IZUMO1-deficient male mice were completely infertile as a result of their sperm cells being unable to fuse with egg cells[6]. It was originally thought that the CD9 protein expressed on the cell surface of oocyte cells was the interacting partner of the IZUMO1 protein[7]. However, IZUMO1 proteins on sperm cells were shown to interact with both wild-type and CD9 deficient egg cells[7], leading to the identification of JUNO as the protein receptor that interacts with the IZUMO1 protein[4]. JUNO is a glycophosphatidylinositol-anchored protein receptor on the cell surface of oocyte cells[4]. The JUNO protein is encoded by the FOLR4 gene, located on chromosome 11 of the human genome[8]. This receptor is a member of the folate receptor family and it was initially referred to as FOLR4. However, since the receptor itself does not actually interact with folate, it was renamed JUNO after the Roman goddess of fertility and marriage[4]. Besides participating in sperm-egg recognition, it has also been suggested that JUNO plays a role in preventing additional sperm fusion with an already fertilized egg[9].

Due to their participation in the early steps of fertilization, it has been suggested that IZUMO1 and JUNO could play a significant role in infertility[10, 11]. The World Health Organization and the Centers for Disease Control and Prevention of the United States have described infertility as a global public health issue, with consequences including psychological distress, social stigmatization, economic strain, and marital discord [12–15]. Globally, infertility affects 15% of couples of reproductive age[16, 17]. In Canada, infertility was estimated in 2011 to affect up to 16% of heterosexual couples, where the woman is age 18 to 44[18]. This statistic has almost doubled since 1992, when 8.5% of women in that age group were considered infertile, and has tripled since 1984, when 5% of women in this age group were considered infertile[18]. Different areas of the world are impacted by infertility at different rates; in 2011, among couples of reproductive age in China, the prevalence of infertility was 25%[19]. In the USA, 6% of married women aged 15 to 44 years were non-surgically infertile[20]. Globally, the age-standardized prevalence rate of female infertility increased by 14.962% from 1990 to 2017[21]. The apparent increase in infertility has led to an increased demand for treatments and diagnosis of infertility. It has been suggested by fertility experts that investigations into the function and mechanisms of JUNO and IZUMO1 could open new doors for diagnosing and treating infertility as well as the design of contraceptives[22].

In this work, the genetic sequences of both IZUMO1 and JUNO protein coding genes in 2,504 humans sequenced in the 1000 Genomes project[23] were retrieved, analyzed and compared. Population-based statistical analysis was used to estimate nucleotide and genetic diversity and deviation from evolutionary neutrality. Haplotype inference and analysis was performed for each gene and across the gene pair. Both genes are located on autosome chromosomes and, therefore, are present in both sexes. However, to identify any selective evolutionary pressure specifically arising from their role in fertilization, we performed most of our analyses for the two sexes together as well separately, within the same population. The results of these analyses were used together to identify signs of selection in these genes. Two large scale scanning methods were also used to independently identify signals of positive and of balancing selection at the chromosome level. In addition, genetically linked missense mutations in IZUMO1 and JUNO were analyzed in the context of protein structures and their function. All the analyses where aimed at determining the degree of sequence conservation among members of the same species, identify signs of co-selection and better understand how genetic diversity plays out on genes that depend on one another for function. The analysis presented here constitutes a starting point for investigators in multiple fields working with IZUMO1 and JUNO, and a potential framework for data analysis of other gene pairs.

## RESULTS AND DISCUSSION

### Dataset

All the analyses presented for human IZUMO1 and JUNO in this manuscript were based on whole-genome sequence data from the phase 3 of the 1000 Genomes project. The 1000 Genomes project dataset corresponds to full-length DNA sequences of 5,008 chromatids from 2504 human individuals belonging to 26 population groups clustered into 5 larger population groups (African (AFR), American (AMR), East Asian (EAS), European (EUR) and South Asian (SAS)). The number of individuals in each population group as well as the composition of the larger groups in terms of the 26 populations are described in Supplementary Materials Table S3. The group labels (geographic and continental) we used in this study follow the original annotations of the 1000 Genomes project. The 1000 Genomes dataset contains variant positions relative to a reference genome (GRCh37) for each of the 5,008 chromatids sequenced. This dataset was also used to calculate Tajima’s D, nucleotide diversity, and overall F_ST_ for selected genes. Our comparisons of Tajima’s D, nucleotide diversity, and overall F_ST_ thus correspond to different genes present **on the same individuals** used in the IZUMO1 and JUNO analyzes. In addition, scanning for signals of selection was performed on the complete sequences of human chromosomes 11 and 19 from the 1000 Genomes project and it is, thus, based on the same samples as the other analyzes. To our knowledge, the 1000 Genomes dataset is one of the most comprehensive large population-based whole-genome dataset publicly available.

### General Statistics

The IZUMO1 and JUNO genetic sequences of 2504 individuals included in the 1000 Genomes project[24] were obtained from the NCBI[25] in variant call format (vcf). The individuals sequenced come from 26 population groups clustered into 5 larger population groups African (AFR), American (AMR), East Asian (EAS), European (EUR) and South Asian (SAS), plus all of the samples (ALL).

SNV density (number of SNVs per 100 bp) in African and non-African population groups for IZUMO1 and JUNO is compared to other genes in Figure 1. We compared IZUMO1 and JUNO to other well-characterized genes to place the frequency at which SNVs are observed in our target genes into a broader context. The data for the reference genes in this analysis was also obtained from the 1000 Genomes project[24] which sequenced the whole genome of the 2504 participants. Therefore this, and all subsequent analysis involving multiple genes, are performed using different genes from the same dataset. The reference genes have been suggested in the literature to be selectively neutral, under balancing selection, or under positive selection from the analysis of different datasets and using different methodologies than the ones in this study (the list of genes used for comparison is found in Table S4).

**Figure 1.**
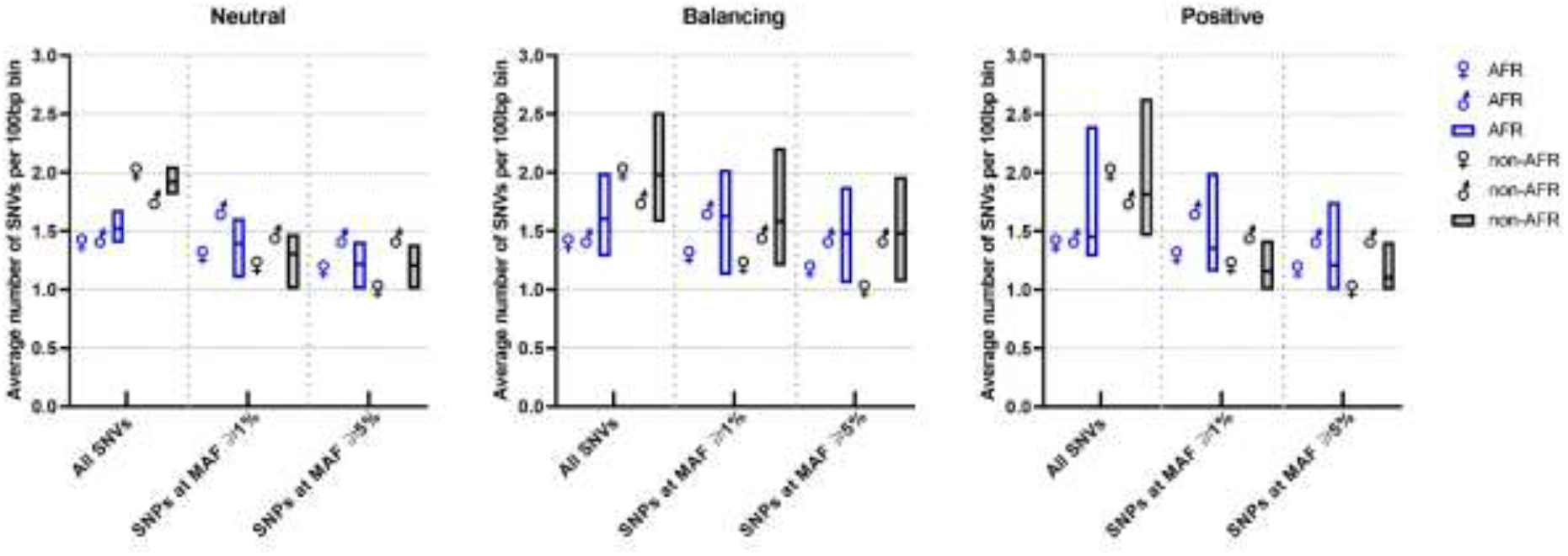
Average number of SNVs per 100bp bin in African (blue) and non-African population (black) groups. Genes previously categorized in the literature as selectively neutral, under balancing selection, or under positive selection. The boxes represent SNV density distribution within each group with the mean marked as a line within the box. IZUMO1 and JUNO are represented by the traditional male and female symbols, respectively. Overall, the total number of SNVs per 100bp bin is higher than the number of SNPs with a MAF ≥1% or ≥5% in non- Africans compared to Africans, indicating a larger number of rare alleles ((MAF < 1%) in non- African populations. In particular, JUNO in non-Africans has a larger number of rare alleles than IZUMO1.

Single nucleotide polymorphisms (SNPs) with minor allele frequency (MAF) ≥1% or ≥5% are distinguished from total SNVs for each group of genes. SNV density is generally higher in non-African populations. This difference can be attributed to a larger number of rare alleles (MAF < 1%) in non-Africans, as indicated by the larger gap between total number of SNVs and number of SNPs at 1% and 5% MAF cutoffs. JUNO in non-Africans has a larger number of rare alleles than IZUMO1. In contrast, IZUMO1 has larger percent of variant sites with a MAF ≥ 1% in both African (56%) and non-African populations (28%) than in the JUNO gene (19% in African and 9% in non-African populations), indicating that JUNO is more conserved across humans than IZUMO1. Neither JUNO or IZUMO1 stand out from the other genes selected for comparison. Genes classified in the literature as under balancing selection trend, on average, towards higher SNV density than genes believed to be under positive selection.

In Table 1, we compare the number of SNVs across 5 populations for both IZUMO1 and JUNO. For both IZUMO1 and JUNO the number of SNVs is decreased sharply when they are filtered with a MAF of 5% or higher, signifying that the majority of SNVs within each of the genes do not appear at significant frequencies throughout the 2504 individuals sampled in the 1000 Genomes Project. The number of SNPs used in linkage disequilibrium (LD) analysis for IZUMO1 was consistent between the AFR, EAS, SAS, AMR and EUR populations. AMR, AFR and EUR males have a higher number of SNPs (1, 2 and 1, respectively) compared to their female counterparts, indicating that the minor allele of these IZUMO1 SNPs appear at a frequency higher than the cutoff, in males but not in females. For JUNO, the number of SNPs used in LD analysis is consistent between most of the grouped populations, except the EUR population. For this population, only one SNP passed both filters for LD analysis, indicating that the JUNO gene is well conserved in EUR population.

**Table 1:** The number of SNPs in IZUMO1 and JUNO in different grouped populations using different filters. The first indicates the total number of SNPs within the entire IZUMO1 and JUNO genes for each population, second row indicates the number of SNPs with a MAF greater than 5% for each population, the third row represents the number of SNPs that meet both 5% MAF and Hardy-Weinberg (HW) p-value, and were used in LD analysis; the fourth row represents the number of SNPs used in LD analysis for just the males in IZUMO1 and just the females in JUNO. The populations analyzed include AMR (n=347), AFR (n=661), EUR (n=503), EAS (504), SAS (489) and ALL (n=2504), where n indicates the number of individuals in each population group.

| Gene | IZUMO1 |  |  |  |  |  | JUNO |  |  |  |  |  |
| --- | --- | --- | --- | --- | --- | --- | --- | --- | --- | --- | --- | --- |
| Population | AMR | AFR | EUR | EAS | SAS | ALL | AMR | AFR | EUR | EAS | SAS | ALL |
| Total Number of SNPs | 70 | 90 | 67 | 60 | 65 | 189 | 24 | 33 | 24 | 21 | 17 | 82 |
| Number of SNPs passing 5% MAF thresholds | 30 | 29 | 30 | 35 | 40 | 31 | 6 | 6 | 6 | 6 | 6 | 6 |
| Number of SNPs passing 5% MAF and HWE threshold for LD analysis- Both sexes | 30 | 29 | 30 | 31 | 31 | 1 | 6 | 6 | 1 | 6 | 6 | 0 |
| Number of SNPs passing 5% MAF and HWE threshold for LD analysis- Males in IZUMO1 and Females in JUNO | 31 | 31 | 31 | 31 | 31 | 2 | 6 | 5 | 1 | 6 | 6 | 1 |

SNPEff[26] indicated that there are 12 (1.7%) missense mutations in IZUMO1. However, only one of them is present with a MAF of 5% or greater in the whole population. In the JUNO gene, the majority of the variants are intronic (49, 56.3%). Only 1 out of 18 missense mutations in the JUNO gene is present with an allele frequency of 5% or greater in the 1000 Genomes phase 3 dataset.

### Tajima’s D test for selection neutrality

Tajima’s D is a null hypothesis test of selection neutrality. Significant deviation from zero indicates that the null hypothesis of neutrality is rejected, and that a non-random process is driving the evolution of the sequence[27]. Generally, a large positive Tajima’s D is indicative of balancing selection, in the absence of a population bottleneck, whereas a large negative value may be indicative of directional selection, linkage between sites or fast population expansion[27]. However, actual values for Tajima’s D depend on the size of the population investigated[28, 29]. The availability of large-scale genome sequencing data has pushed the traditional population size limits from 100’s to well above 1000’s, requiring contextual adjustments. Moreover, the magnitude of this size effect is modulated by the type of segregating sites (e.g., neutral versus constrained) included in the calculation[28]. To place the Tajima’s D values we calculated for IZUMO1 and JUNO into context, we selected 39 genes known to be under selection neutrality, positive, or balancing selection and calculated Tajima’s D values for these genes using the same procedures and gene sequences from the same individual as IZUMO1 and JUNO (Figure 2). In each case, Tajima’s D was calculated using all biallelic sites within the gene segment, averaged over 100 bp bins, for all populations combined (n=2504 individuals), the AFR population (n=661) or the combined non-AFR populations (n=1843) sequenced in the 1000 Genomes project. As can be seen in Figure 2, the general trend is for Tajima’s D values to be higher for genes under balancing selection than for genes under positive selection, with neutral genes between the two. IZUMO1 clusters with balancing selection genes in both the African versus non-African distribution graph as well as in the all- populations normalized means. In contrast, JUNO clusters with the genes under positive selection. The 23 genes known to be under positive selection have fairly negative Tajima’s D, with an all-populations mean value of -0.71±0.02, consistent with an excess of rare alleles. In contrast, the 12 genes under balancing selection have a much wider spread of Tajima’s D values, ranging from -0.58 to 0.66, with a negative mean value (−0.17 ± 0.12). Genes under balancing selection are generally expected to have large positive Tajima’s D in the absence of a population bottleneck. A possible reason for this discrepancy is the inclusion in our calculations of rare alleles that may not have been included in the studies that originally classified some of the genes in the balancing selection category. As an example of this effect, if only sites with a MAF of 1% or above are used in Tajima’s D calculation (as opposed to also including all rare sites), the mean value for genes under balancing selection increases to 1.37 ± 0.34, a positive value statistically above 0 (two-tailed p value < 0.0001) as expected. Although a more comprehensive review of reference genes is in order, the results shown in Figure 2 give a clear estimation of where IZUMO1 and JUNO lie with respect to genes previously suggested to be under selection in the literature: IZUMO1 Tajima’s D indicates balancing selection, whereas JUNO’s Tajima’s D is consistent with positive selection. The Tajima’s D values for the reference genes are reported in Supplementary Table S4, along with some literature values[30–37].

**Figure 2:**
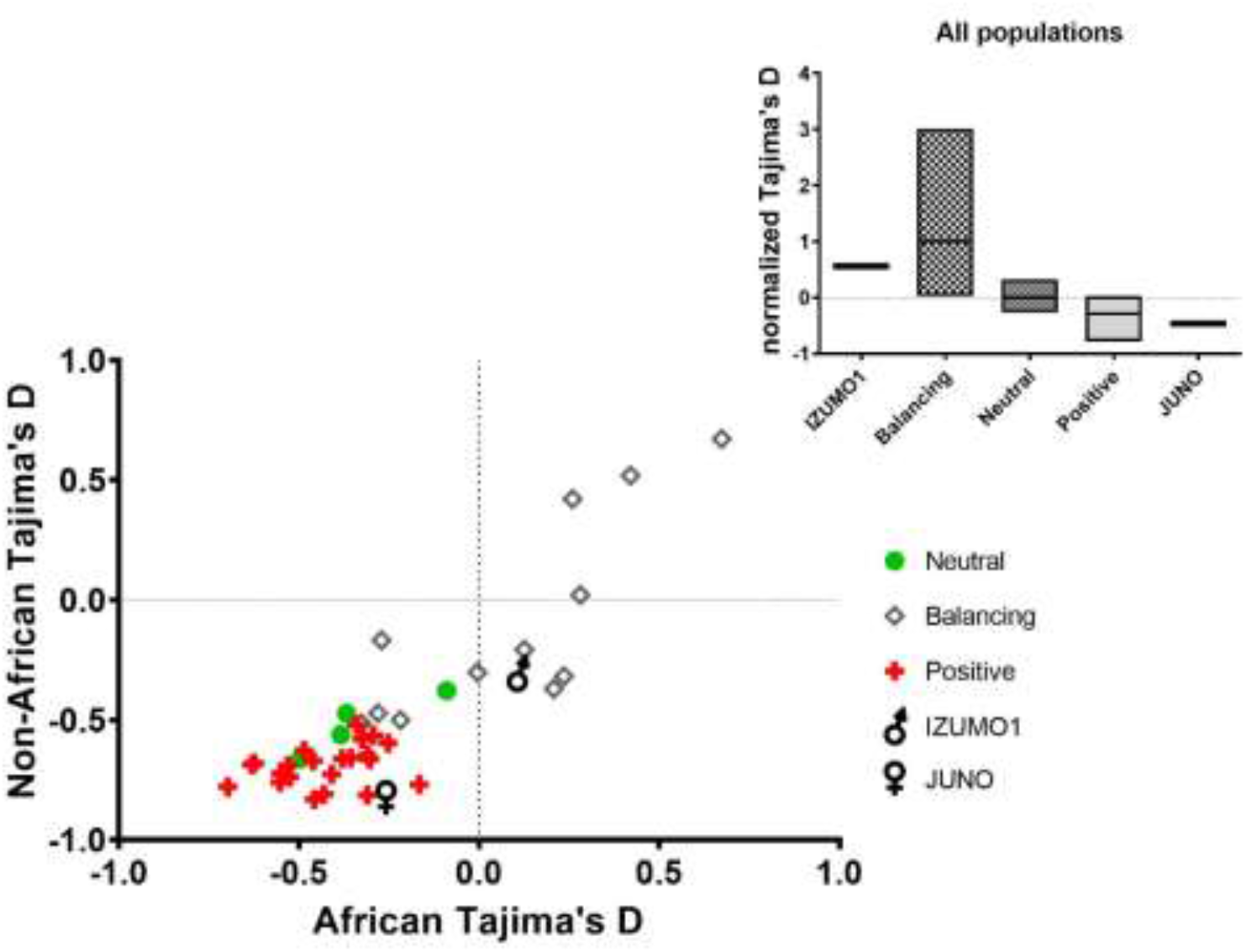
A comparison of the Tajima’s D values of neutral genes (circles) and genes believed to be under balancing (diamonds), or positive (crosses) selection. IZUMO1 and JUNO are represented by the traditional male and female symbols, respectively. The mean (bar) values for all populations combined are shown in the insert within maximum and minimum value boxes.

### Nucleotide diversity

Nucleotide diversity (π) was estimated for the same genes and populations used for Tajima’s D as an additional characterization of selection (Figure 3). Nucleotide diversity is expected to be low for loci under positive selection in comparison neutral loci or to loci under balancing selection. In Figure 3, the genes assumed to be under positive selection have lower average π value than neutral genes or the genes under balancing selection, with the average π value for neutral genes falling between the positive and balancing selection groups. IZUMO1’s average π is higher than the maximum value within the neutral gene and the positive selection groups, suggesting its placement in the balancing selection group. JUNO’s average π value, on the other hand, is comparable to the average π value for the positive selection group and much lower than the balancing selection or the neutral groups, supporting its placement in the positive selection group.

**Figure 3:**
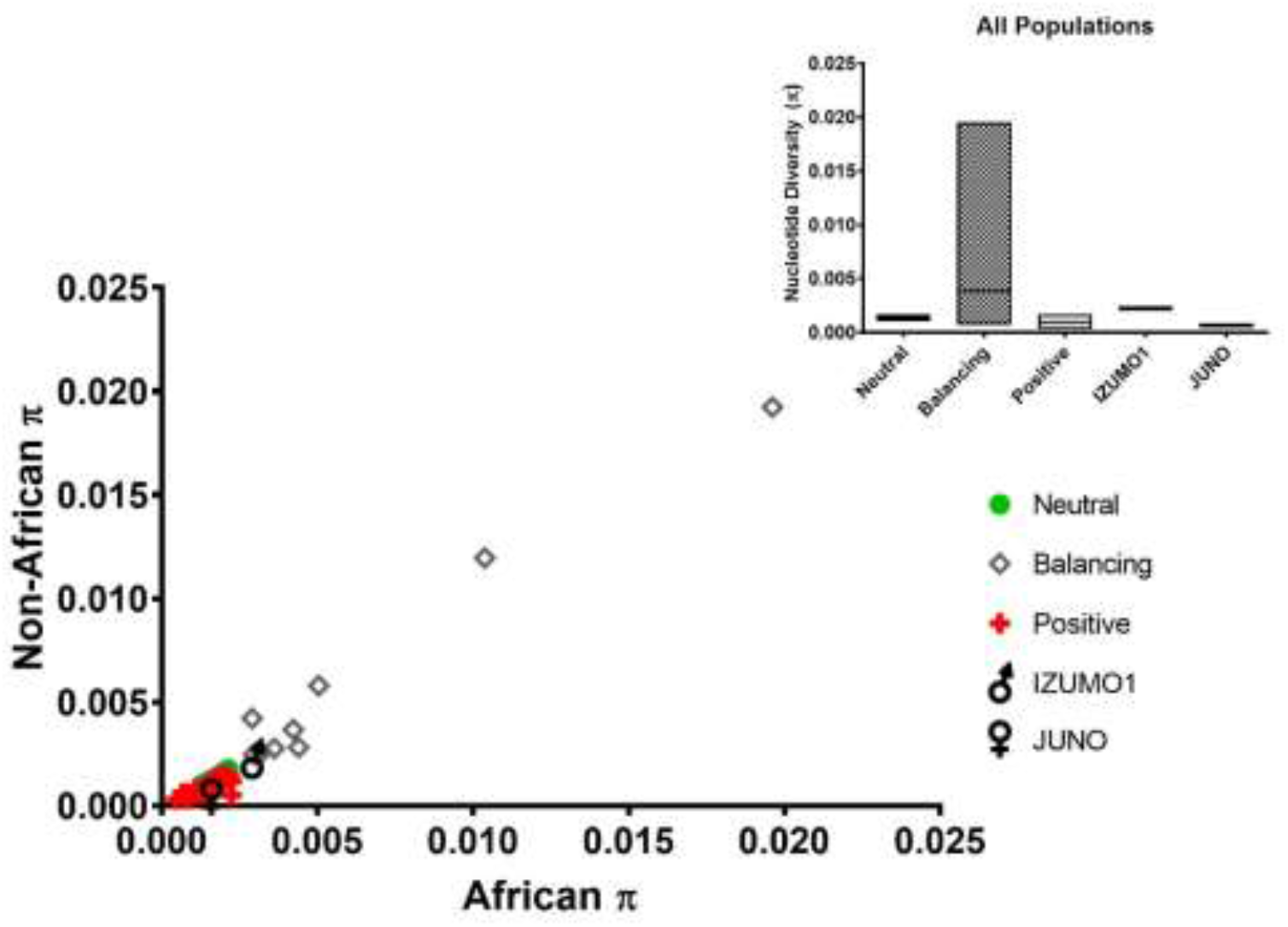
A comparison of the nucleotide diversity (π) values of genes believed to be under, selection neutrality (circles), balancing (diamonds) or positive (crosses) selection. IZUMO1 and JUNO are represented by the traditional male and female symbols, respectively. The mean (bar) values for all populations combined are shown in the insert within maximum and minimum value boxes.

### Haplotype inference

Haploview[38] was used for haplotype inference. The dataset used in this analysis is the same one used in all t\he other analyzes presented so far. Haploview selects variant sites for LD analysis and haplotype inference based on two criteria: minor allele frequency (MAF) and Hardy-Weinberg Equilibrium (HWE). In our haplotype analysis, only biallelic sites with MAF≥ 5% and HWE p-value > 0.01 were considered. Interestingly, while most variant sites present HWE p-values that differ significantly from zero (p-value > 0.01) and can, thus, be considered in equilibrium (i.e., HWE assumptions are satisfied) for each gene in both sexes together or separately, some exceptions were observed when individuals in EAS and SAS populations were grouped into a single Asian population group (ASI) for IZUMO1 (data available in Supplementary materials Table S5). These deviations also affect the ALL group which includes all population groups together. Although the ASI supergroup presented variant positions that are not in Hardy-Weinberg equilibrium, the individual groups EAS and SAS have all loci in Hardy-Weinberg equilibrium. The HWE p-values of the ASI supergroup suggest population stratification, wherein there is a systematic difference in allele frequencies between subpopulations in a population, possibly due to different ancestry. For this reason, EAS and SAS are treated as separate groups in our population-based analyses. In another intriguing observation, when comparing the HWE p-value of the JUNO loci in both sexes and in just females (Supplementary materials Table S6) we see some sex-based differences in the AFR and AMR populations in all 5 positions examined (rs61742524, rs55784852, rs16920146, rs7925833, and rs7935583). The females in these populations present a much lower chance probability than both sexes combined, with most positions falling from ∼50% to less than 18%, and some becoming as low as 2%. The difference between the observed and predicted heterozygosity in the JUNO gene of AFR and AMR females could be partially associated to a violation of one of the HWE rules. Loci not in HW equilibrium were excluded from LD calculation and haplotype inference. The number of variant sites passing MAF and HWE p-value thresholds are presented in Table 1 for both IZUMO1 and JUNO genes.

A haplotype analysis was carried out for IZUMO1 and JUNO using the variant sites passing MAF and HWE thresholds. The individual sequences were grouped according to sex (male and female), and larger groups (AMR, AFR, EUR, EAS, SAS) of the original 26 populations. In addition, population analysis was performed using sequences from only females for the JUNO gene and from only males for the IZUMO1 gene, since the role of these proteins in reproduction can only be fully understood in the gametes of their respective sex. The data collected from this analysis is presented in Tables 2 and 3.

**Table 2:** The haplotypes and their haplotype frequencies in the IZUMO1 gene (gene ID 284359) for both sexes analyzed together as well as separately in different grouped populations. The underlined nucleotide indicates a missense polymorphism where G is part of a codon for Ala and A is part of a codon for Val at amino acid position 333 in the protein. The locations outlined are rs2307018, rs2307019 and rs838148 in this order. These locations had r^2^ values greater than 0.007 and D’ values all greater than 0.998. The populations analyzed include AMR (n=347), AFR (n=661), EUR (n=503), EAS (504), SAS (489) and ALL (n=2504), where n indicates the number of individuals in each population group. It should be noted that the IZUMO1 gene is found in the reverse strand.

| Population | IZUMO1 Haplotype Frequency |  |  |  |  |  |  |  |  |
| --- | --- | --- | --- | --- | --- | --- | --- | --- | --- |
|  | <u>CGA</u> |  |  | <u>AAG</u> |  |  | <u>CGG</u> |  |  |
|  | All | Males | Females | All | Males | Females | All | Males | Females |
| AFR | 0.35 | 0.36 | 0.34 | 0.32 | 0.31 | 0.32 | 0.33 | 0.32 | 0.33 |
| AMR | 0.16 | 0.17 | 0.16 | 0.54 | 0.54 | 0.55 | 0.29 | 0.29 | 0.29 |
| EUR | 0.09 | 0.08 | 0.10 | 0.45 | 0.43 | 0.46 | 0.46 | 0.48 | 0.44 |
| EAS | 0.14 | 0.15 | 0.16 | 0.83 | 0.84 | 0.82 | 0.01 | 0.01 | 0.01 |
| SAS | 0.29 | 0.30 | 0.27 | 0.50 | 0.48 | 0.52 | 0.22 | 0.22 | 0.21 |
| ALL | 0.22 | 0.22 | 0.22 | 0.51 | 0.51 | 0.52 | 0.27 | 0.26 | 0.26 |

**Table 3:**
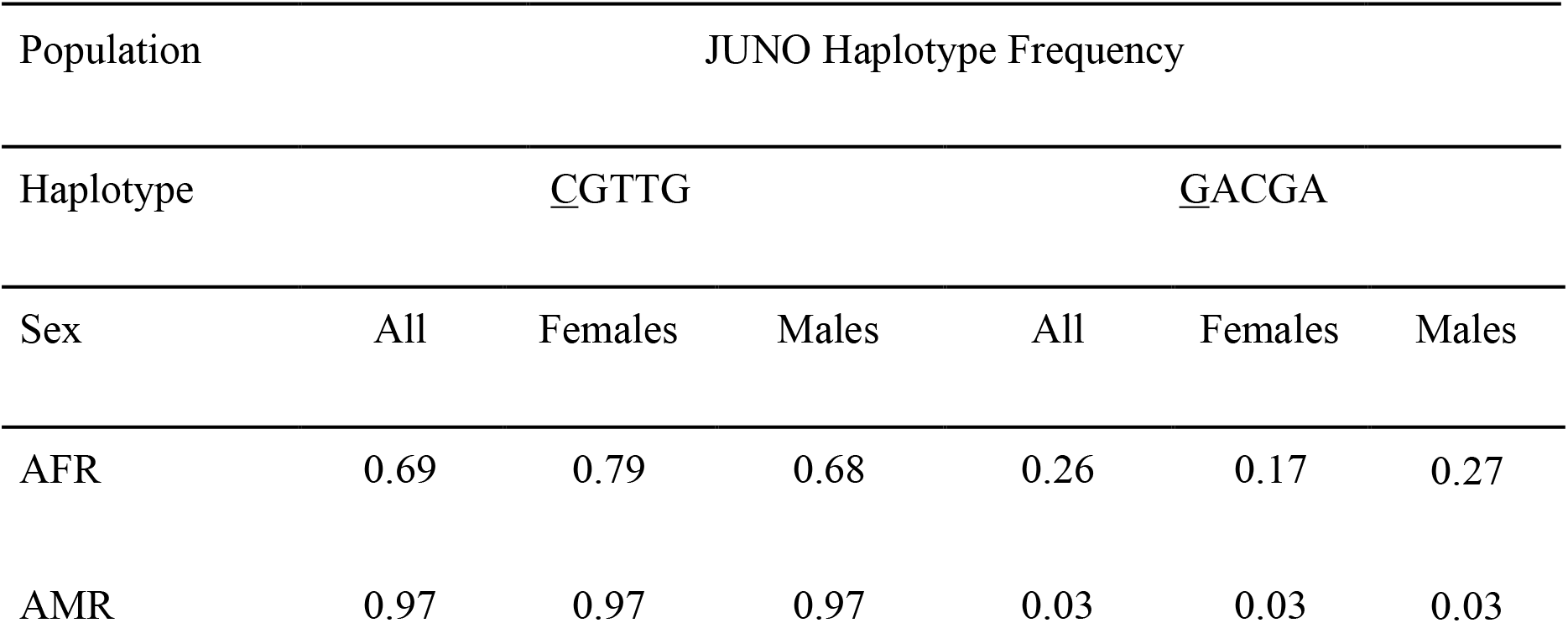
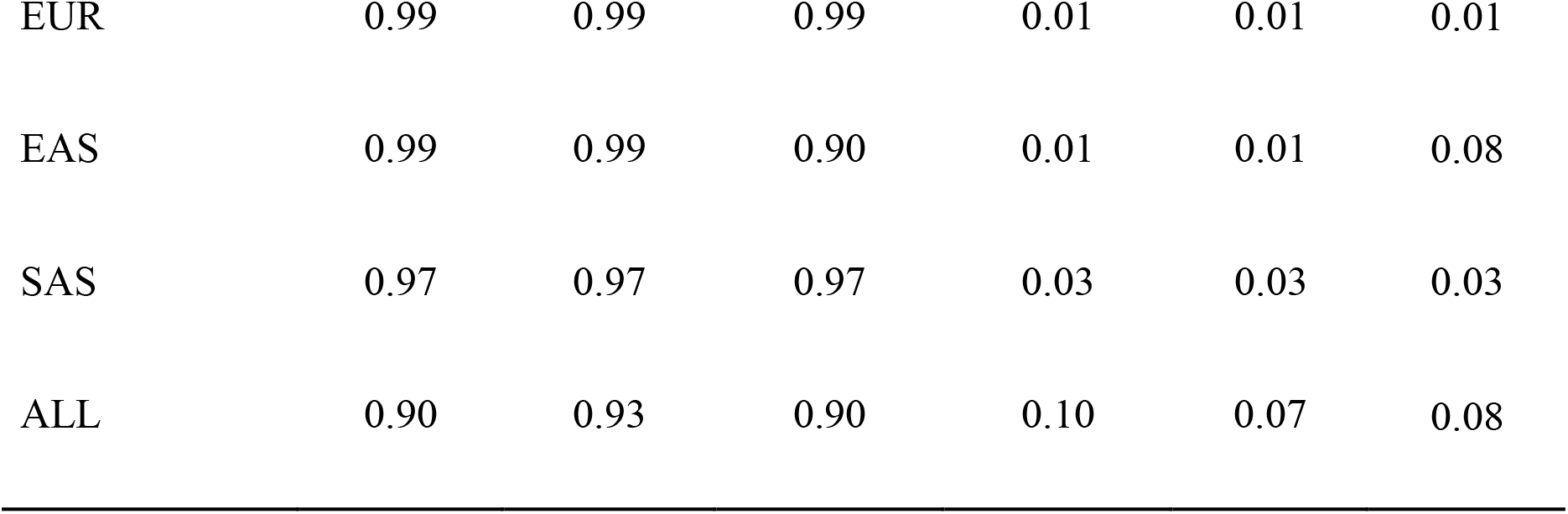
The haplotypes and their haplotype frequencies in the JUNO gene (gene ID 390243) for both sexes analyzed together as well as separately in different population groups. The underlined nucleotide indicates a missense polymorphism at rs61742524 where C is part of a codon for Cys and G is part of a codon for Trp at amino acid position 3 in the protein. The locations outlined are rs61742524, rs55784852, rs16920146, rs7925833 and rs7935583 in this order. The r^2^ values were all greater than 0.851 and D’ values are all greater than 0.997. The populations analyzed include AMR (n=347), AFR (n=661), EUR (n=503), EAS (504), SAS (489) and ALL (n=2504), where n indicates the number of individuals in each population group.

Analysis of linkage disequilibrium can be heavily influenced by sample size and it is generally believed that relatively large samples are required for detection of disequilibrium[39–41] . How large of a sample is needed to detect LD with high confidence (>90%) depends on the frequencies of the alleles under LD investigation[41]. Based on a study published by Thompson et al. [41], we can estimate the sample size needed to determine LD with 90% confidence by considering the minor allele frequencies of the SNPs under investigation. For IZUMO1 (rs2307018, rs2307019 and rs838148) the minor allele frequencies range from 0.4 (40%) to 0.2 (20%), whereas for JUNO (rs61742524, rs55784852, rs16920146, rs7925833 and rs7935583) the minor allele frequencies are all around 0.1 (10%). Investigating positive LD for any 2 loci in the MAF range of 0.1 to 0.4 with 90% confidence using a test size of 0.05 requires samples ranging in size from 14 to 68 individuals, depending on the frequencies of the pair of loci. The smallest population group collected from the 1000 Genomes Project is 347 individuals in the AMR group, a number over 5 times higher than the required sample size suggested by Thompson et. al., giving a high level of confidence in our results.

The haplotypes reported all contain missense mutations, which are underlined in the haplotypes in Tables 2 and 3. The missense mutation in the IZUMO1 sequence is a conserved substitution (Ala to Val), not expected to dramatically impact protein structure and function. For JUNO, the 2 amino acids substituted in the missense mutation are characteristically different: Trp is an aromatic amino acid with a hydrophobic character, whereas Cys has a polar neutral side chain. This amino acid substitution could affect protein function.

For IZUMO1, there are 3 haplotypes that appear equally frequently in the AFR population, with no sex difference. Differences between 1% and 2% in the haplotype frequency of males only compared to both sexes and females only are seen in the other populations. However, these differences could be attributed to incorrect haplogroup assignment due to error in variant calling, which is estimated to be between 0.1% and 1%, incorrectly[42, 43]. EAS, AMR and SAS populations all have 1 haplotype that is dominant (≥50% of the population), A<u>A</u>G. A secondary high-frequency haplotype is present in the AMR and SAS population groups at the same frequency, 29%, but with different allele combinations (C<u>G</u>G and C<u>G</u>A, respectively). Interestingly, EAS presents overwhelmingly as A<u>A</u>G (83%), whereas SAS is 54% A<u>A</u>G and 29% C<u>G</u>G indicating that these two populations do not form a homogeneous group together (they are often grouped as “Asian” in the literature). The haplotypes A<u>A</u>G (45%) and C<u>G</u>G (46%) are evenly distributed in the EUR population group. The presence of multiple haplotypes at considerable frequencies in IZUMO1 is consistent with balancing selection. Two of the three variant positions that are part of the haplotypes for IZUMO1 are found in NCBI’s SNP database (dbSNP)[44], however no information on health-related associations are available. dbSNP indicates that rs2307019 is a complex locus which appears to produce several proteins with no sequence overlap. This locus is shared by two different genes, IZUMO1 and RASIP1[44]. In the RASIP1 gene the SNV rs2287922 converts Arg to Cys[44]. This is not a conserved substitution changing from a large, positively charged, basic side chain (Arg) to a medium-sized, polar, uncharged side chain (Cys), and therefore could impact protein function. The RASIP1 gene encodes for the RASIP1 protein, which is required for the proper formation of vascular structures that develop both vasculogenesis and angiogenesis; it acts as a critical and vascular-specific regulator of GTPase signalling, cell architecture, and adhesion, which is essential for endothelial cell morphogenesis and blood vessel tubulogensis [45]. It should be noted that the large MAF seen for this SNP (rs2307019) could be due to its possible effects on the regulation of RASIP1 expression.

For JUNO, all population groups present one dominant haplotype (<u>C</u>GTTG) with frequency at or above 97%, except AFR. A 9% sex bias is seen in AFR for this haplotype. Sex bias is not observed in any other population group. AFR shows more diversity than the other population groups with the second haplotype present in 26% of the population. A third haplotype that is only present in the AFR population was identified but at a low frequency (<u>G</u>ACTA, 6%). This haplotype is not observed in any other population group. This further supports the conclusion that the largest difference in the JUNO gene is seen in the AFR population. Both the r^2^ values and the D’ values for the JUNO haplotypes are high, indicating strong linkage between all the loci involved in the haplotypes. The dominance of one haplotype in JUNO is consistent with both selection neutrality and positive selections, which may be distinguished from one another by determining the ancestral alleles which we will discuss in later sections. All five positions in the haplotype appear in dbSNP[44]. There are no known clinical significance associated with most of these positions. However, the missense mutation at position rs61742524 (c.9C>G, p.Cys3Trp) may be clinically relevant. This mutation was found in 3 of 330 females with polyspermy but not in any of the 300 matching controls in a study involving *in-vitro* fertilization[46]. The haplotype carrying this mutation, <u>G</u>ACGA, is significantly present in the AFR population (26% frequency) and, interestingly, displays a sex bias with a lower prevalence in females (17%) compared to the group as a whole (26%). The lower frequency of this haplotype in AFR females may reflect impaired fertility in females carrying this mutation. The frequency of this allele in AFR is much higher than in the other populations, suggesting the existence of a genetic advantage that has yet to be identified for the carriers of the G allele. Further investigation of the role of JUNO in biological processes other than reproduction may provide insights on the evolutionary role of the <u>G</u>ACGA JUNO haplotype.

### Frequencies of combined haplotypes

IZUMO1 and JUNO are a receptor-ligand pair encoded in 2 autosome chromosomes. Due to their role in fertilization, their expression and function is well-documented in gametes but not in other cell types. IZUMO1 is overwhelmingly expressed in testes, with low levels of expression in lungs, stomach, and adrenal glands, among other organs[47, 48]. JUNO presents high expression levels in ovaries and adrenal glands, and it is also expressed in lymph nodes, spleen, and thyroid, among other organs[47, 48]. It is reasonable to expect that this receptor-ligand pair interact with one another in other organs outside their fertilization role and, if so, the 2 genes may be evolutionary linked albeit not by physical proximity. To explore this possibility, we first looked for intergenic LD but none was found between pairs of SNPs with MAF ≥ 1% (the highest r^2^ for any intergenic pair of SNPs was 0.0193) . To test for indirect co-dependency between the 2 genes, we calculated the frequencies at which each JUNO haplotype is observed in combination to each IZUMO1 haplotype within the same chromatid. This data is shown in Figure 4. The expected frequencies considering random combination of haplotypes from each gene is also presented in Table 4. If the two genes were genetically linked their observed combined haplotype frequencies would differ from the frequencies expected as a result of random combinations. As observed in Table 4 and Figure 4, there is a slight preference for the IZUMO1-JUNO haplotype combination A<u>A</u>G-<u>C</u>GTTG over A<u>A</u>G-<u>G</u>ACGA, likely due to the female bias found for <u>C</u>GTTG in AFR. However this difference is not significant, as determined by Wilcoxon matched-pairs test yielding a two-tailed p-value of 0.99. Overall, we find no indication of strong genetic linkage between the two genes, since their combined haplotypes are observed at the same frequencies they are expected to be observed by chance.

**Figure 4.**
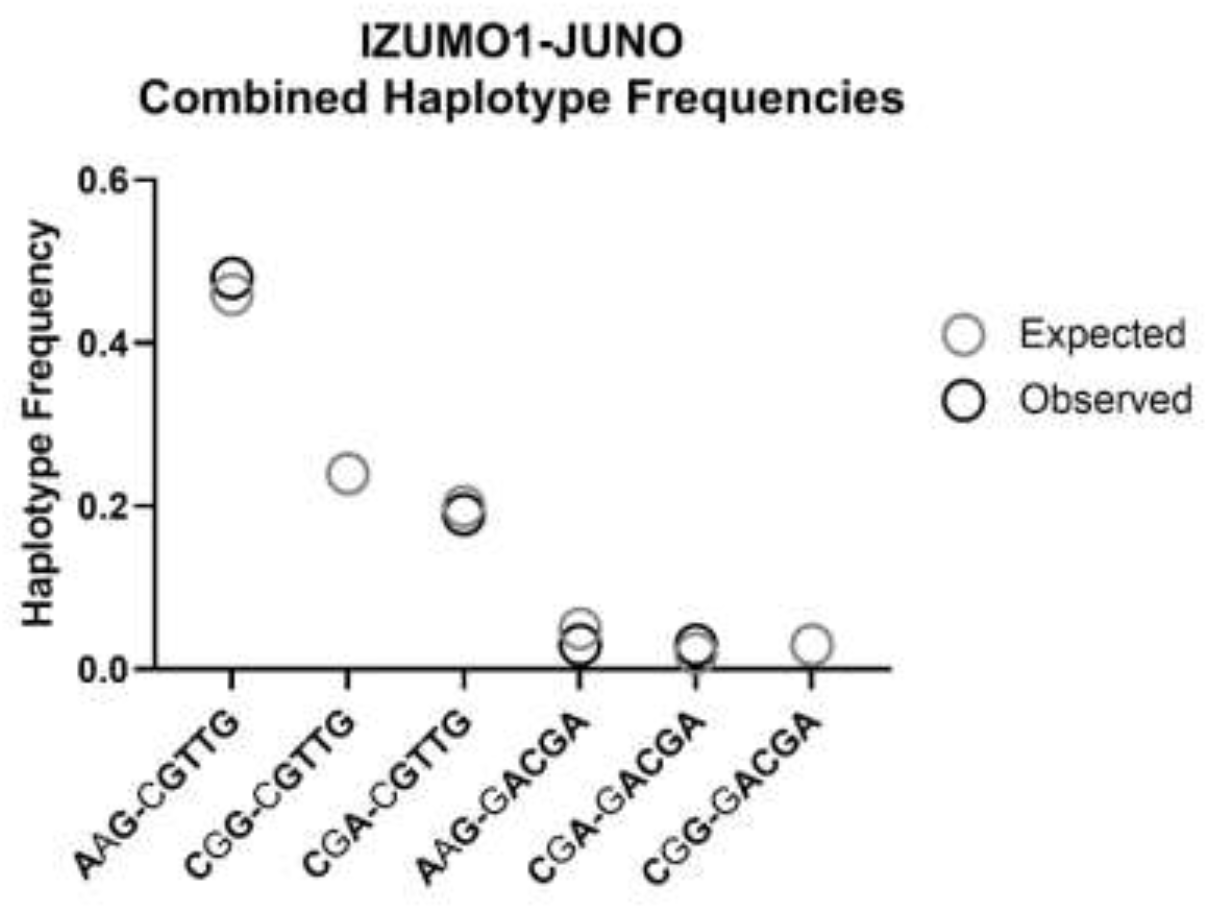
Expected (grey circle) versus observed (black circle) frequencies for combined IZUMO1 and JUNO haplotypes.

**Table 4:**
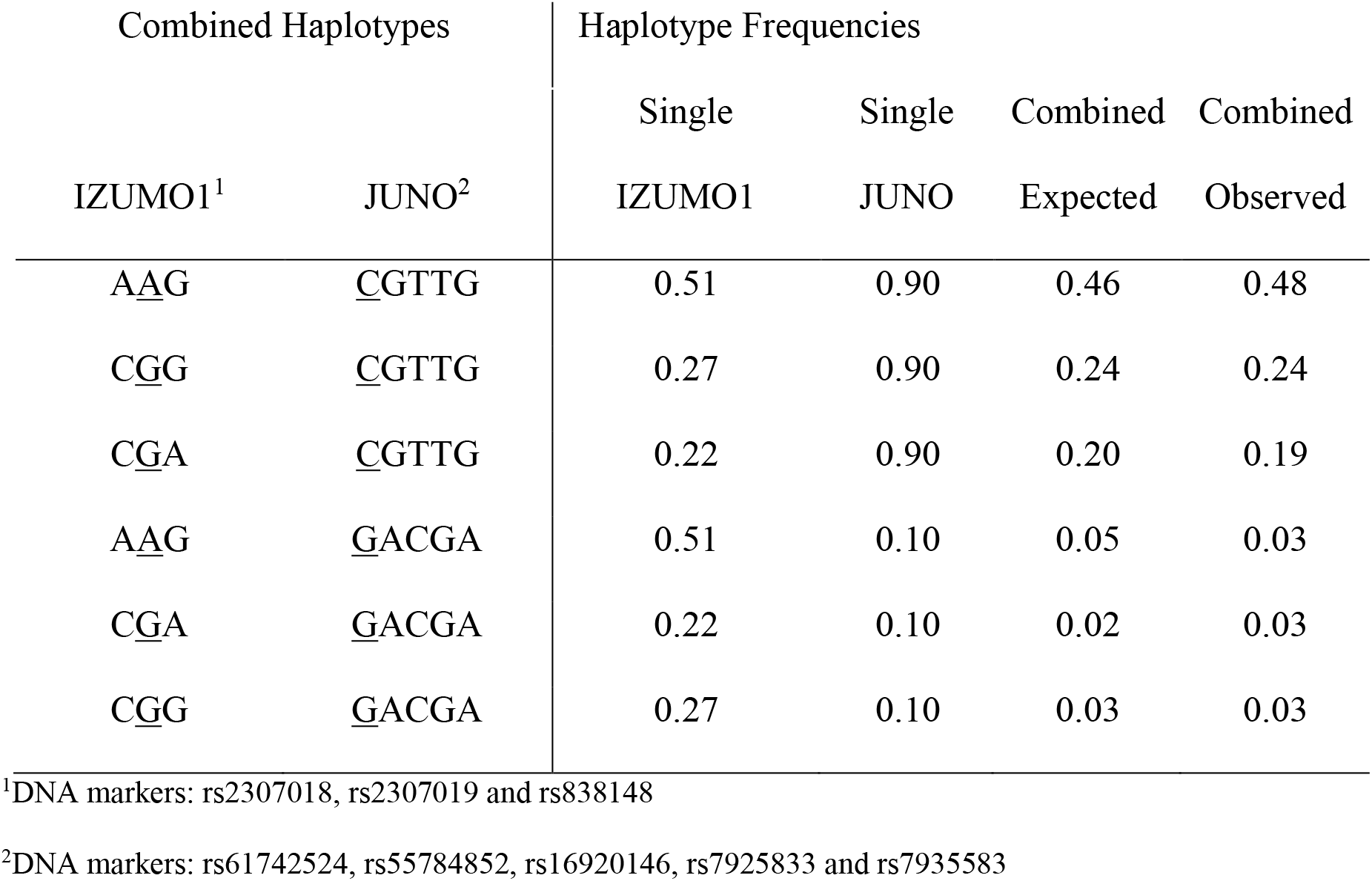
Frequency of combined haplotypes for IZUMO1 and JUNO genes. Haplotype frequencies for each individual gene are provided under “Single” columns. The frequency expected if assortment of cross-gene haplotypes is random (“Expected”) is compared to the frequency of both haplotypes appearing together as observed in the phased gene sequences analyzed (“Observed”). Underlined nucleotides correspond to missense mutations.

#### Structural Analysis of Haplotypes

The structures of the IZUMO1 and JUNO proteins, as well as their complex have been determined experimentally[49–52]. On its own, IZUMO1 adopts a distinct boomerang shape with a rod-shaped N-terminal, four helix bundle (4HB), C-terminal immunoglobin-like domain (Ig-like domain) and two anti-parallel beta strands that function like a hinge between the 4HB and Ig-like domain [50–52]. JUNO contains five short alpha-helices, three 3_10_ helices, and two short two-stranded antiparallel beta sheets, stabilized by eight conserved disulfide bonds[50–52]. JUNO consists of a rigid core and three surface-exposed, flexible regions[52].

These proteins bind with high affinity in a one-to-one ratio via the N-terminal of IZUMO1, resulting in major conformational changes observed in the four-helix bundle (4HB) and hinge region of IZUMO1. IZUMO1 abandons its boomerang shape and adopts a stabilized and locked upright conformation[51] upon binding. It has been proposed that Trp62 in JUNO interacts with hydrophobic residues in the alpha-helical core (4HB) of IZUMO1 (residues 57-113)[52]. Through mutational studies, it has been concluded that the IZUMO1-JUNO interface is resilient to mutations because it is stabilized through the combined effects of multiple van der Waals, hydrophobic, aromatic and electronic interactions[51].

Neither one of the non-synonymous variant positions that are part of our haplotypes for IZUMO1 and JUNO is found in the experimentally determined structures. The experimentally determined structures of IZUMO1 normally span from amino acids 22 to 255 (out of 355 amino acids), which represent the extracellular regions of the protein. The non-synonymous SNP in our haplotype (rs2307019) encodes for position 333 of the cytoplasmic domain of IZUMO1 protein, which is located outside the boundaries of the currently available 3D structures. This position it is not located near the interaction surface of these two proteins (residues 57-113) [52] and, therefore, the Ala333 to Val substitution in the IZUMO1 protein is unlikely to have an impact on the binding of IZUMO1 and JUNO proteins. However, this missense mutation is also an upstream modifier of the RAIP1 protein, which could explain its high MAF. Two other variant positions (rs2307018 and rs8108468) in IZUMO1 with a MAF of at least 5% are both synonymous variants. One of them, rs2307018, is also an upstream modifier for the RASIP1 protein. The other synonymous variant, rs8108468, is also an upstream modifier of the RASIP1 protein as well as a downfield modifier of the FUT1 protein. This restriction may act as an evolutionary pressure on RASIP1 and/or FUT1 and further investigation, although outside the scope of this work, is warranted.

The currently available experimentally determined structures of JUNO span from amino acid 20 to 228, encompassing the majority of the 250 amino acids of this protein. The SNP of interest in our haplotype (rs61742524) is part of the codon position 3 of the JUNO protein which is not included in any of the currently available experimentally determined structures. The fact that the missense mutation (Cys3Trp) occurs far away from the interaction surface of the two proteins (roughly from Lys42 to Tyr 147)[50] makes it unlikely to adversely impact their binding ability. However, the potential role of this SNP (rs61742524) in polyspermy is interesting; after fertilization, there is normally a rapid loss of JUNO from the oolemma by cleavage, which is a possible mechanism for blocking polyspermy in mammals[46]. JUNO is anchored to the membrane at position 228 on the C-terminal of the protein which is tagged with a GPI moiety[53]. Although position 3, which is the one modified by polymorphisms on rs61742524, is far from the anchoring position, the processes leading to membrane anchoring and later cleavage involves interactions with multiple proteins[54], some of which may be impacted by amino acids substitutions at the N-terminal of JUNO. There has been also suggestions in the literature that the amino acid substitution at position 3 affects the expression of JUNO at either the mRNA or the protein level [46].

Due to the lack of experimentally determined structures with the mutations highlighted in this work (Ala333Val in IZUMO1 and Cys3Trp in JUNO) an additional approach was used to assess the effect of these substitutions. PolyPhen-2, a human nonsynonymous SNP prediction database (Polymorphism Phenotyping v2; http://genetics.bwh.harvard.edu/pph2/)[55], was employed to analyzed the resulting amino acid sequences and predict the possible impacts of the 2 amino acid substitutions on the structure and function of IZUMO1 and JUNO. PolyPhen-2 is an automatic tool for predicting the possible impact of amino acid substitutions on the structure and function of a human protein based on the sequence, phylogenetic and structural, information characterizing the substitution[55]. The Ala333Val mutation in the IZUMO1 protein was determined to be “probably damaging” with a score of 1.00. The Cys3Trp mutation in the JUNO protein was determined to be “benign” with a score of 0.00[55].

### Differentiation between population groups

The Weir-Cockerham fixation index (F_ST_) values between different populations were calculated. F_ST_ is a comparative measure of the average number of pairwise differences between individuals sampled from different population groups and the average number of pairwise differences between individuals sampled from the same population group[56]. We selected as a reference the average F_ST_ value of 0.124. This number is an average of 3 human genome wide F_ST_ values reported in the literature. The first (0.123) was calculated for the entire human genome using 26,530 autosomal SNPs from 3 different populations (African-American, East Asian and European-American)[57]. The second genome- wide F_ST_ value (0.119) was calculated by Cavalli et al. using 120 “classical” non-DNA polymorphisms in 42 worldwide populations[58]. A third study that analyzed 8,525 autosomal SNPs in 84 African-American, European-American, Chinese and Japanese individuals described an F_ST_ value of 0.130[59]. These values represent genetic diversity across human populations as reflected throughout their genomes, and we consider their average of 0.124 ± 0.006 to be an appropriate benchmark to which we compare our own data. To further support this choice of benchmark value, we calculated overall F_ST_ for the set of reference genes used in the Tajima’s D and nucleotide diversity analysis, and found the mean of these values (0.103 ± 0.063) to be consistent with the genome-wide literature-based value but carrying a much larger standard deviation. Thus, an F_ST_ value for the IZUMO1 and JUNO genetic sequences of the different population pairs that is equal or lower than 0.124 plus or minus two standard deviations is deemed to be consistent with genome wide F_ST_ values and in line with the amount of variation seen within the entire human genome. The pairwise F_ST_ values for each pair of population groups for IZUMO1 and for JUNO are shown in Figure 5 (the complete set of F_ST_ values between all populations are displayed in Supplementary Tables S7 and S8). For IZUMO1, EAS (gray circles) is the only population group that seem to diverge from the genome-wide value with an average F_ST_ of 0.291 ± 0112. However, this divergence is short of statistical significance (p-value = 0.06 in a two-tailed t-test), indicating that the differentiation seen in this gene between all four continental populations and EAS is still in line with the differentiation seen within the entire human genome. The average F_ST_ value (0.266 ± 0.051) between all four continental populations for JUNO with respect to AFR is significantly higher than the genome wide F_ST_ average (p-value = 0.01 in a two-tailed t-test), indicating a significant degree of population divergence. This divergence likely results from a 26% prevalence of the JUNO haplotype <u>G</u>ACGA in the AFR population, but only 3% or less prevalence of this haplotype in the EUR, SAS, AMR, and EAS population groups. The average F_ST_ values for EUR, SAS, AMR, and EAS population groups in the JUNO gene are in line with the benchmark genome-wide value.

**Figure 5:**
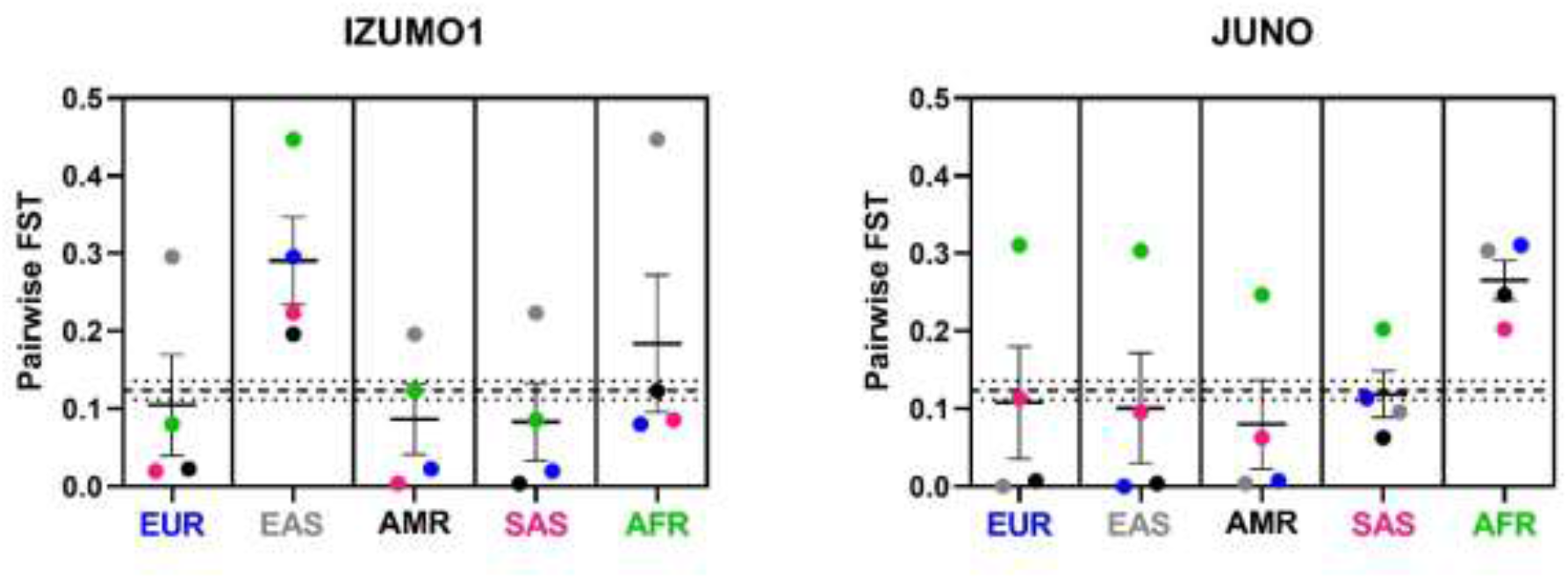
F_ST_ values in the IZUMO1 and JUNO genes for the European (EUR), East Asian (EAS), South Asian (SAS), African (AFR) and American (AMR) population pairs. These F_ST_ values were calculated using SNPs with a MAF of at least 5%. An F_ST_ along the 0.124 (dashed line) plus or minus 2 standard deviations (dotted lines) is considered to be consistent with the rest of the human genome[57]. The average F_ST_ value for each population group relative to all other groups is shown as a solid line. The populations analyzed include AMR (n=347), AFR (n=661), EUR (n=503), EAS (n=504), and SAS (n=489), where n indicates the number of individuals in each population group.

The pairwise F_ST_ values of population groups for all individuals, and for just males or just females with respect to EAS for IZUMO1 and to AFR for JUNO are presented in Figure 6. In this figure, the values are with respect to the population group we found to have the highest degree of genetic diversity compared to others (Figure 5 analysis). Thus, the graphs in Figure 6 represent the most substantial pairwise F_ST_ values in the data sets.

**Figure 6:**
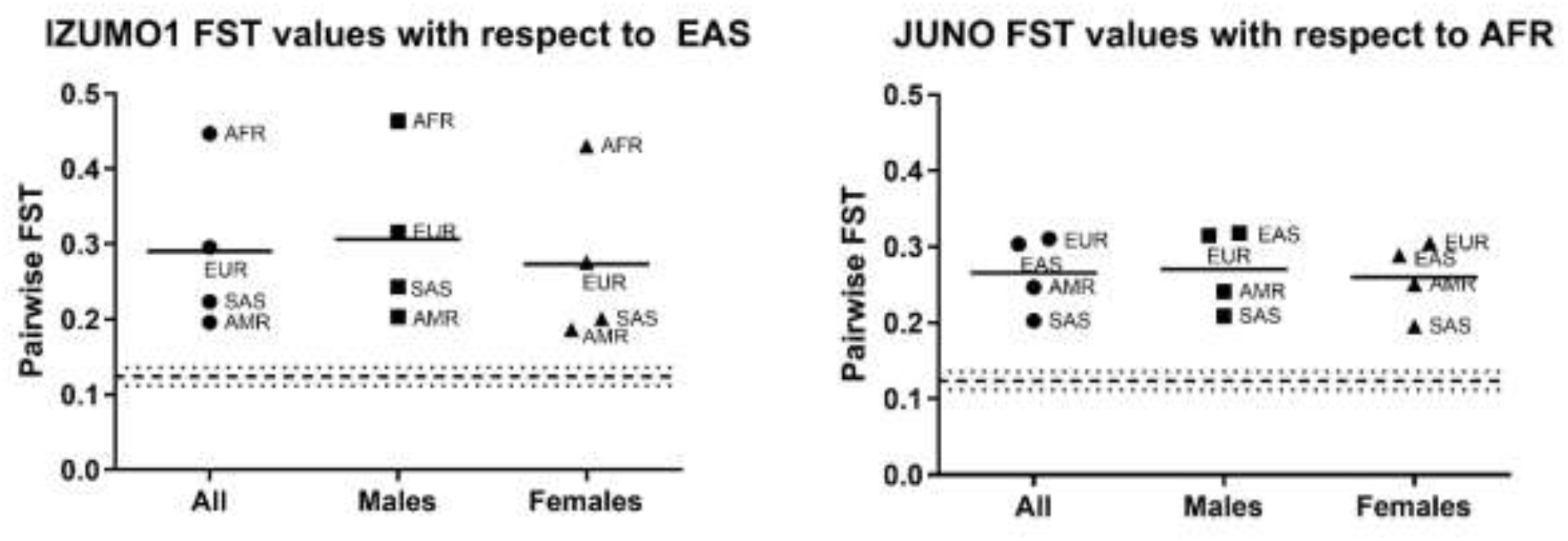
F_ST_ values for the IZUMO1 gene with respect to the East Asian (EAS) and for the JUNO genes with respect to the African (AFR) populations. Each population group is analyzed in its entirety as well as for each sex. These F_ST_ values were calculated using SNPs with a MAF of at least 5%. An F_ST_ along the 0.124 value (dashed line plus or minus dotted lines) is considered to be consistent with whole human genome[57]. The average F_ST_ value (solid line) for each sex is consistent with the average F_ST_ value for both sexes combined. The populations analyzed include AMR (n=347), AFR (n=661), EUR (n=503), EAS (n=504), and SAS (n=489), where n indicates the number of individuals in each population group.

F_ST_ between the males and the females of all four of the continental populations was 0.0008 for IZUMO1 and 0.0007 for JUNO, respectively. Both are close to zero, indicating that there are no significant differences between the female and male sequences of IZUMO1 and JUNO. These F_ST_ values were calculated using SNPs with a MAF of 1%.

To further investigate population diversity, we calculated pairwise F_ST_ values associated to all 26 original populations that form the 5 continental groups EAS, EUR, SAS, AMR and AFR, considering only SNPs with a MAF of 5% when all individuals from all populations are grouped together. We used these values in a principal component analysis (PCA) to map genetic diversity between the 26 population groups represented in the 1000 Genomes dataset (Figures 7 and 8). PCA was used as a way to better visualize pairwise F_ST_ values and, thus, understand population diversity. Population based F_ST_ values are traditionally compared relative to one of the populations and depict how all the others differ from that one population, as opposed to one population to another. Although overall F_ST_ can be used to compare population diversity, two populations with very distinct F_ST_ profiles could still have similar overall F_ST_. The advantage of PCA analysis is that it combines pairwise F_ST_ for each population and displays their variance in a “map” where proximity among populations indicates similarity in diversity profiles, as measured by pairwise F_ST_. We find this representation more informative, as it compares the populations to each other in terms of their diversity profiles, as opposed to an average diversity index or a comparison to a reference population.

**Figure 7:**
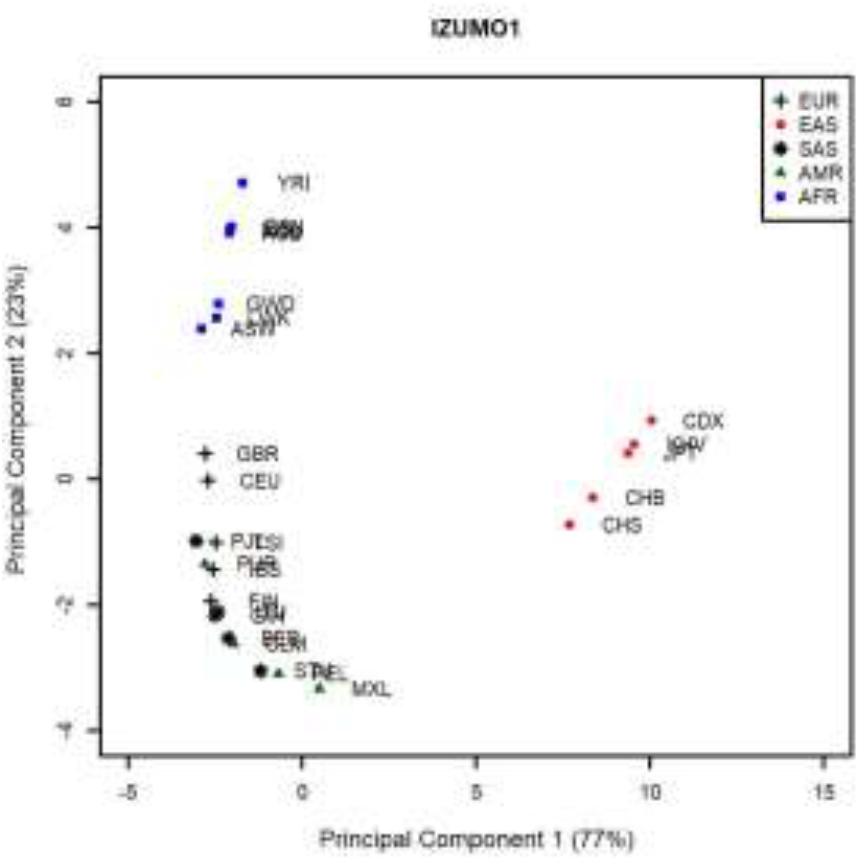
Principal component analysis of the F_ST_ values for the IZUMO1 gene between 26 population groups that constitute the supergroups AFR (square), EUR (cross), EAS (circle), AMR (triangle), and SAS (star). The percentages between parentheses correspond to the variance carried by each PC. EAS and AFR segregate from the other 3 population groups.

**Figure 8:**
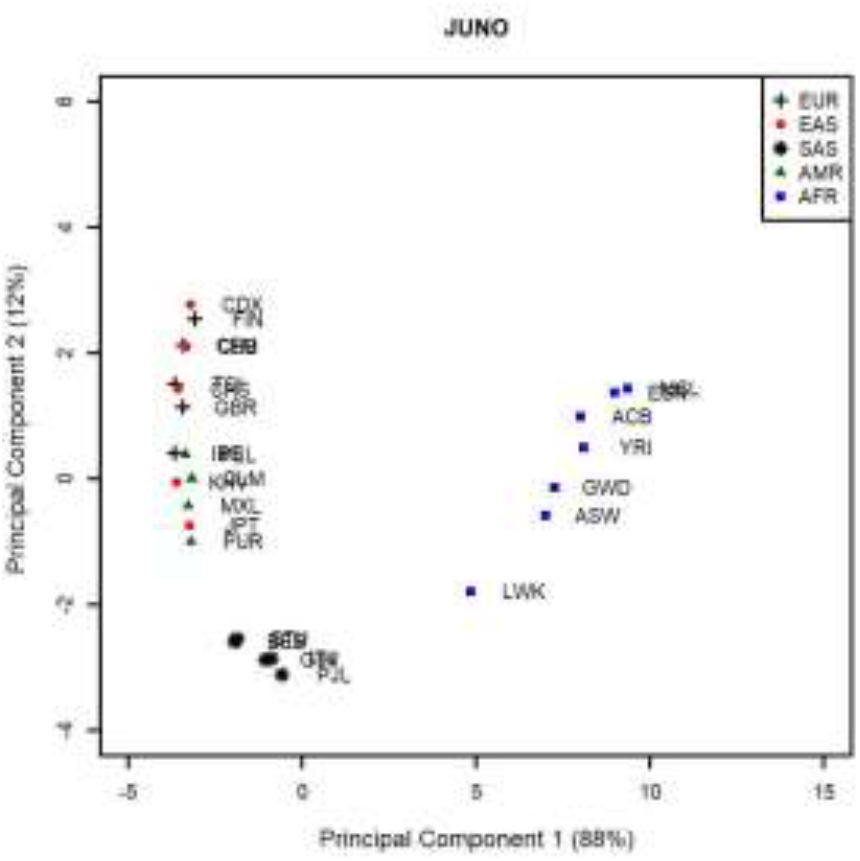
Principal component analysis of the F_ST_ values for the JUNO gene between 26 population groups that form the supergroups AFR (square), EUR (cross), EAS (circle), AMR (triangle), and SAS (star). The percentages between parentheses correspond to the variance carried by each PC. AFR and SAS segregate from the other 3 population groups.

Core geographic populations belonging to the group AFR (squares) cluster together and are separate from the other core populations for both IZUMO1 and JUNO. Populations belonging to the EAS group (circles) are separate from the others for IZUMO1, but overlap with populations belonging to the EUR (crosses) and SAS (stars) groups for JUNO. Similarly, core geographic populations in the AMR group (triangles) are separate for JUNO but overlap with other geographic groups for IZUMO1. For IZUMO1, the core populations GBR and CEU segregate from the other EUR groups and do not overlap with any other core populations. Similarly, the core population LWK (AFR) segregates from the other AFR core populations for JUNO. These results do not support a model of co-selection between IZUMO1 and JUNO, as the pattern of genetic diversity represented by the PCs is not as similar as it would have been expected if these 2 genes were co-selected.

### Haplotypes Present in Archaic Genomes

Genomes of archaic members of the genus *Homo* were downloaded from the Max Planck Institute for Evolutionary Anthropology (http://cdna.eva.mpg.de/neandertal/altai/) in vcf format, and analyzed to identify alleles at locations corresponding to haplotypes for IZUMO1 (Table 2) and JUNO (Table 3) in contemporary humans.

Two out of 3 DNA markers for IZUMO1 (rs2307018 and rs2307019) are present in the genome sequences of the Neanderthal from the Vindija Cave[60] and the 45,000 year old anatomically modern human from Ust’-Ishim[61]. Ust’-Ishim is heterozygous at both positions, whereas Vindija is a homozygous C<u>G</u>. A C at rs2307018 and G at rs2307019 correspond to either a C<u>G</u>A or a C<u>G</u>G haplotype in Table 2. The frequency of the C<u>G</u>G haplotype is most prominent in EUR (0.46). C<u>G</u>A is present at moderate frequencies in AFR (0.35) and SAS (0.29). Assuming the same pattern of heritance, the heterozygous Ust’-Ishim likely had one A<u>A</u>G parent. This haplotype is predominant in EAS (0.83) and AMR (0.54), and is prominently present in EUR (0.45).

JUNO’s DNA markers rs61742524, rs55784852, rs16920146, rs7925833 and rs7935583 were present in the genomes of the Vindija cave Neanderthal[60], Altai Neanderthal, and Denisova. All 3 genomes have the haplotype <u>C</u>GTTG, which is the major haplotype in present-day humans analyzed in this report. This haplotype has a frequency greater than 0.97 in every population except the AFR population (0.69). This could indicate that the JUNO gene has diverged from its ancient genetic sequence through selective pressures in AFR, or that inter-breeding between modern humans, Neanderthal and Denisova brought this haplotype to higher frequencies in all but AFR populations. As we shall see in the next section, the later is the most likely to have happened.

### Haplotype present in other primates

A pre-built genomic alignment of 37 mammals[62] was used to identify the ancestral allele in each locus used in our haplotype inference. We used the flanking sequences reported for each loci in the dbSNP database[44] to correctly identify each allele within the alignment, instead of relying in position numbering. For the IZUMO1 the ancient alleles at positions rs2307018, rs2307019, and rs838148 are CGG in this order. This allele combination corresponds to haplotype C<u>G</u>G in our analysis which is 1 of 2 equally represented haplotypes in EUR (0.46 frequency) and it is 1 of 3 equally represented haplotypes in AFR (0.33 frequency). The dominant haplotype in EAS, A<u>A</u>G (0.83 frequency), contains 2 derived alleles out of 3, whereas the third IZUMO1 haplotype (C<u>G</u>A) contains one derived allele at the last position. The A<u>A</u>G haplotype is well-represented in all 5 populations, albeit it is only dominant in EAS, and may have resulted from a founder effect or a selective sweep in this population. For JUNO, the ancient alleles at positions rs61742524, rs55784852, rs16920146, rs7925833 and rs7935583 are GGCTG, in this order. This allele combination does not correspond to either one of our haplotypes. The haplotype (<u>C</u>GTTG) that we identified as prevalent in all of the 5 populations analyzed here contain 2 derived alleles, whereas the second JUNO haplotype (<u>G</u>ACGA), which is mostly present in AFR populations, contains 3 derived alleles. The existence of multiple high frequency derived alleles within a relatively small genome segment is consistent with positive selection[37].

### Interspecific Sequence Comparisons

There are many homologous mammalian species for IZUMO1 and JUNO protein sequences. However, only 29 mammalian species with available matches to both the *Homo sapiens* IZUMO1 and JUNO amino acid sequences were used in this study. These sequences were aligned with ClustalX[63]. The percent identities for each of the sequences in reference to the *Homo sapiens* amino acid sequences for both JUNO and IZUMO1 can be found in Supplementary Table S9 and S10. With the highest percent identities being in the high 90s and the lowest percent identities being around 50% for IZUMO1 and 70% for JUNO. The sequence for the IZUMO1 protein is less conserved than that of the JUNO protein, which is indicated by the low percent identities seen between the homologous species in reference to the human sequence.

### Intraspecific Sequence Comparisons

The nucleotide sequences of all the 2504 individuals from the 1000 Genomes Project for JUNO and IZUMO1 were aligned using ClustalX[63]. The percent identity values shown in Supplementary Table S11 indicate that the JUNO genetic sequences are highly conserved within the human species, with an average percent identity of 99.9%. IZUMO1 genetic sequences are slightly less conserved, with an average percent identity of 99.5%. In comparison, lowland gorilla JUNO and IZUMO1 genetic sequences are 99% and 98% identical to the corresponding human sequences, respectively.

### Scanning for signals of positive selection at the chromosome level

We used RAiSD[64] to scan for signals of positive selection on the DNA sequences of human chromosome 11 (where the JUNO gene is located) from the same 2504 individuals whose IZUMO1 and JUNO sequences are the subject of our study. The total number of SNPs analyzed in this set is 3,865,304 and all 2504 individuals sequenced were considered in the µ statistics calculations as a single group (Supplementary Materials Figure S5), as well as separate population groups (Figure 9). Signals of positive selection correspond to µ values above the threshold (solid red line) which was set to be above the 99.95 percentile of µ scores across the entire chromosome, for each population. These scores are calculated over sliding windows. Straight green dashed lines are used to mark the limits of the JUNO gene in the figure. As figure 9 indicates, for each of the 5 population groups the genomic region corresponding to JUNO is at the edge of a region of high µ values. This region corresponds to the highest peak in the µ value landscape of chromosome 11. The location of JUNO suggests that the indicators of positive selection we identified from the analysis of nucleotide diversity, Tajima’s D, and haplotype inference could have originated from a hitchhiking effect of a selective sweep [65], as opposed to JUNO itself being the target of positive selection. Further investigation is necessary to identify gene(s) within the region of high µ values are direct target(s) of positive selection. However, this outside the scope of the present study.

**Figure 9.**
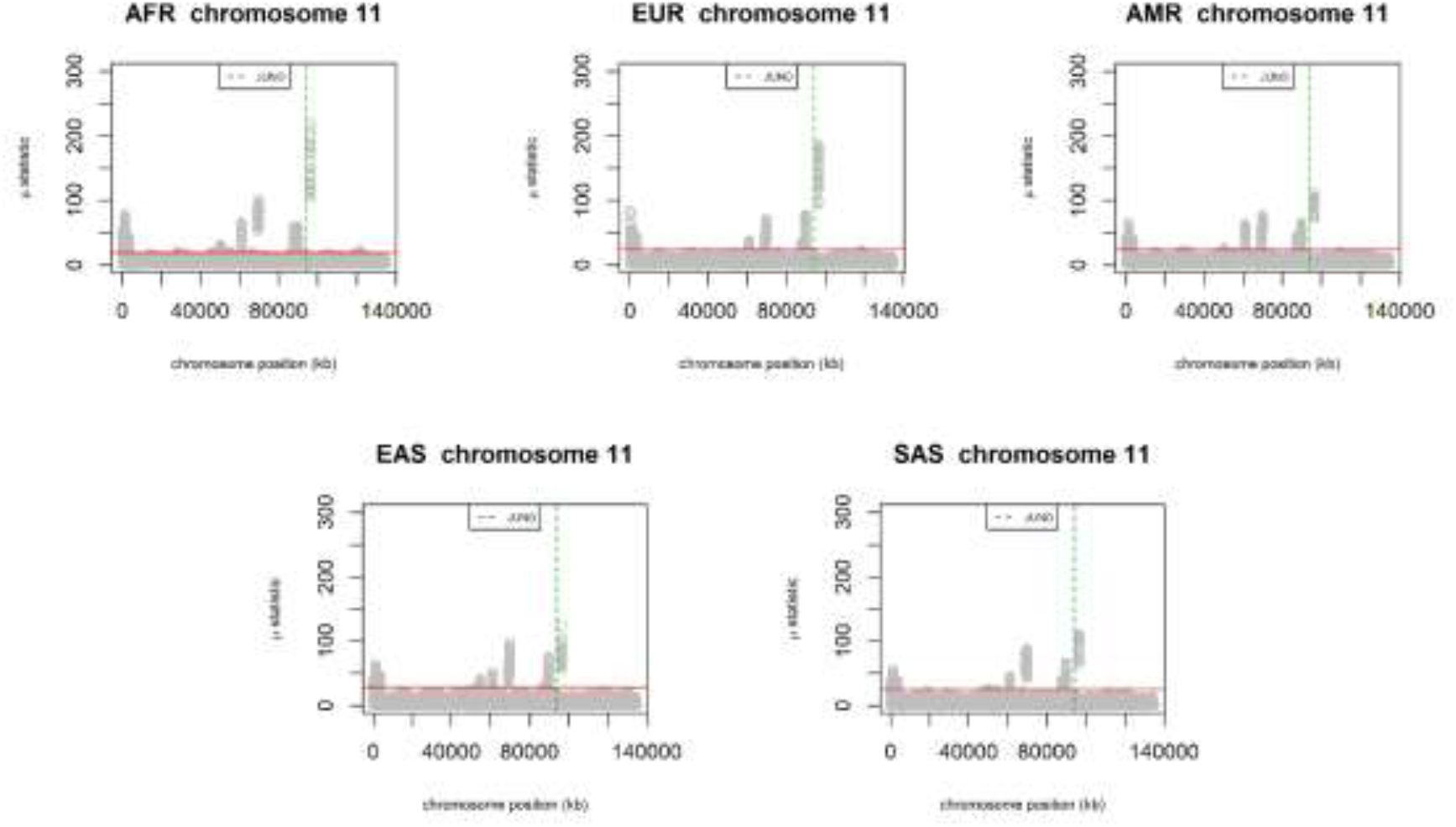
Scanning for signals of positive selection on chromosome 11 of individuals sequenced in the 1000 Genomes project belonging to the AFR, EUR, AMR, EAS, and SAS populations. Regions with µ scores above the 99.95% threshold (solid red line) are expected to be under positive selection. The genomic region corresponding to the JUNO gene is marked by green dashed lines.

### Scanning for signals of balancing selection at the chromosome level

As an independent verification that IZUMO1 is under balancing selection, we scanned chromosome 19 (where IZUMO1 is located) for signals of balancing selection using BetaScan2[66]. This computer application identifies signals of balancing selection using both polymorphism and substitution data. The tool calculates β statistics on a sliding window around each SNP in the dataset. Regions under balancing selection display higher positive beta scores compared to neutral regions. The entire set of 1,746,975 SNPs of chromosome 19 from the 1000 genomes project was used in this analysis. Due to size restrictions of the file converter tool glactools[67], this analysis has to be limited to specific populations. This is in contrast with RAiSD which can handle all 5,008 chromatid sequences in the 1000 Genomes dataset. In our preceding analysis EUR (404 individuals) and AFR (602 individuals) have haplotype frequency distributions consistent with balancing selection, in addition to other metrics that are also indicative of balancing selection. Assuming our assessment of IZUMO1 is accurate then we expect β values associated to IZUMO1’s genomic region to be in the top 99.90 percentile of β scores (solid red line in the figure). This is what we see in Figure 10, which depicts the results of the balancing selection scan of chromosome 19 for the AFR, EUR, AMR, EAS, and SAS populations. The β values associated to IZUMO1’s genomic region are well above the threshold for all 5 populations, and this region displays high values relative to other peak regions in the β statistics landscape of chromosome 19.

**Figure 10.**
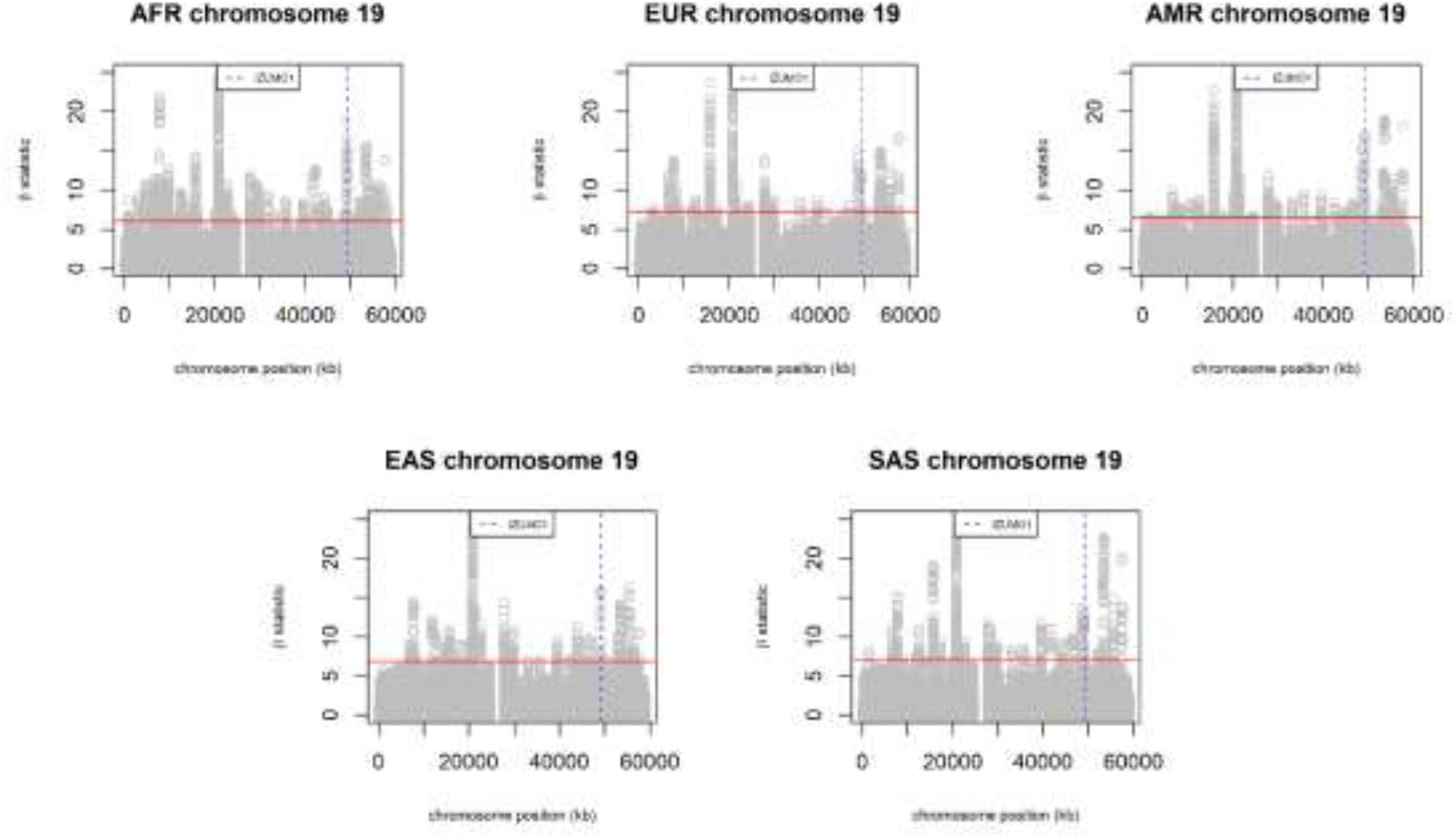
Scanning for signals of balancing selection om chromosome 19 of the AFR EUR, AMR, EAS, and SAS populations of the 1000 Genomes project. Regions with β scores above the 99.9% threshold (solid red line) are expected to be under balancing selection. The genomic region corresponding to the IZUMO1 gene is marked by blue dashed lines.

## CONCLUSION

The IZUMO1 and JUNO genes encode 2 protein partners that play an essential role in fertilization. In this manuscript we present the first large population genetics study on the IZUMO1 and JUNO genes, with some very interesting results. It was expected that the IZUMO1 and JUNO genes would be well-conserved between human populations, given their essential role in fertilization. In general, this held true with both genes demonstrating pairwise-F_ST_ values below a genome-wide F_ST_ average obtained from multiple studies. However, there were some pairwise F_ST_ values that indicated greater diversity between populations, such as all other continental populations with respect to EAS in the IZUMO1 gene, and with respect to AFR in the JUNO gene. The trend associated to the African population group in the JUNO gene is consistent with positive selection which, as we will discuss later, is supported by other metrics.

Interestingly, a JUNO haplotype, <u>G</u>ACGA, that has very low frequency in all other populations has a frequency almost ten-fold higher in the AFR population, suggesting that a selective advantage that has yet to be identified is carried by this haplotype. All 3 archaic genomes we examined have the haplotype <u>C</u>GTTG, which is the major haplotype in present-day humans, and primates carry the C allele in the first position of this haplotype. This indicates that the G allele at locus rs61742524 is a derived allele. Since the G allele is associated to polyspermy, which is disadvantageous for female reproduction, its relatively high frequency in AFR is likely associated to a non-reproductive role of JUNO. Investigating the role of the JUNO protein in non-reproductive cells may, thus, shed light on functional effects associated to the rs61742524 G allele. We found no LD between intergenic alleles and no significant differences between observed and predicted combined IZUMO1-JUNO haplotype frequencies indicating that the two genes are not genetically linked within the same genome. Thus, cross-genes haplotype analysis failed to provide evidence of co-selection for this pair of interacting genes, and so did PCA genetic diversity mapping.

Metrics such as SNV density, nucleotide diversity (π) and Tajima’s D, as well as haplotype analysis and derived allele determination, all support the notion that the IZUMO1 gene in humans is under balancing selection whereas the JUNO gene is under positive selection. These assertions are further supported by chromosome-level scanning for signals of positive and balancing selection on chromosomes 11 and 19, respectively. The indication that IZUMO1 is under balancing selection seems to contradict that in mammals, many male reproductive genes have been shown to evolve rapidly through positive selection[68–70] On the other hand, a study conducted by Grayson and Civetta investigated positive selection of IZUMO genes in mammals and in this study they did not find evidence of positive selection in Glires or Primates in the IZUMO1 gene[71].

Regarding the impact of genetic diversity on fertility, a study using recombinant proteins showed that hamster JUNO can directly interact with mouse, pig and human IZUMO1, matching the ability of zona-free hamster eggs to fuse with human, mouse and pig spermatozoa[72]. This indicates that the differentiation between IZUMO1 and JUNO genes in these homologous species is not great enough to inhibit the recognition step of fertilization. According to our sequence comparison, the percent of identical residues between the human IZUMO1 gene sequence and the mouse sequence (the most distant homolog in the list) is 47%, whereas the lowest percent identity for the human JUNO gene in reference to 29 homologous species is 66% between the mouse and human sequences. Since the sequences of both IZUMO1 and JUNO are above 99.5% identical among human individuals, one should assume that there is not enough diversity in IZUMO1 and JUNO gene sequences among humans to inhibit the recognition step of fertilization in any significant way at the population level. However, rare mutations (≤ 1% MAF) affecting either protein at key interaction amino acids could negatively impact fertility on an individual basis. Rare mutations affecting expression levels of either protein could also negatively impact fertility.

Even though these genes code proteins that directly interact in a key step of fertilization. A great deal of tolerance in diversity of the IZUMO1 sequence, matched by a great deal of conservation of the JUNO sequence, seems to be sufficient to establish some degree of genetic “checks and balance” for this pair. The high degree of conservation combined to a dominant haplotype across multiple population groups makes the protein encoded by JUNO a potential target for the development of contraceptive treatments (with the caveat that such treatments would have to pass the zona pellucida barrier). These contraceptive treatments could include small molecules designed to bind to the JUNO protein in the region of the binding interactions to the IZUMO1 protein, delivered systemically or locally.

Currently, there is a demand for population-based analysis workflows due to the increasing availability of full genome sequences. This demand is compounded by the realization that, in order to fully understand how genomes work, we must extend traditional population analysis to an interacting network of genes context. This study attends that demand by providing a framework for comprehensive analysis of multiple interacting genes.

## MATERIALS AND METHODS

There are a wide variety of population genetic approaches to analyze human genes, however, in this work, only a few were selected to best answer the question of how inter-dependent the IZUMO1 and JUNO are among human populations. A method based on population allele frequency differences was selected (pairwise F_ST_) to investigate the genetic diversity seen in the IZUMO1 and JUNO genes between human populations[73]. We also selected a method based on allele frequency spectrum within each population (Tajima’s D)[74] to investigate selective pressures acting upon each of the genes. Additionally, the Hardy-Weinberg Equilibrium p-values, linkage disequilibrium and haplotype frequencies were inferred using Haploview[38] and utilized to determine patterns in frequencies that would indicate selective pressures acting upon the genes as well as any suggestion of complementary haplotypes in IZUMO1 and JUNO. These population genetics methods have been used in other works to investigate selection and therefore, are appropriate tools for us to use in our investigation of IZUMO1 and JUNO [33, 75–78]. In addition, we used whole-genome scanning methods to identify signals of positive (RAiSD version 2.9 [64]) or balancing (BetaScan2[66]) selection on human chromosomes 11 and 19. Unless otherwise noted, data was analyzed either through applications installed on Lakehead University’s Galaxy server (http://wesley.lakeheadu.ca:8800), which is only accessible within the University’s network, or through in-house developed R[79] Scripts utilizing a variety of R packages.

### Data sources

Data from phase 3 of the 1000 Genome Project[23] was retrieved in variant call format (vcf) from (https://www.ncbi.nlm.nih.gov/variation/tools/1000genomes). We obtained vcf files for IZUMO1 and JUNO as well as 40 selected genes used as reference, and for chromosomes 11 and 19. All of these are from the same set of 2504 individuals whose genomes were fully sequenced by the 1000 Genomes Project. The genomic locations (chromosome:first-last genomic coordinate) for IZUMO1 and JUNO were, respectively, 19:49244073-49250166 and 11:94038803-94040858. The genomic locations of the other genes are reported in Table S4 of Supplementary Material. A detailed analysis of the variant sites in each of the genes is included in Supplementary Tables S1 and S2. A comprehensive description of the individuals sampled is presented in Supplementary Table S3. The number of individuals in each continental population were as follows: South Asian (n= 489), East Asian (n=504), European (n= 503), American (n=347) and African (n=661). The 1000 Genomes project was created in 2008 and provides a freely accessible public database of all the genomes the project has sequenced[23]. In addition to nucleotide sequence information about each individual sequenced, the project also provides information about where they are from, their sex and their relatedness to other individuals sequenced[23]. VCF files simply include the mismatched nucleotides of the sequence for each individual in reference to the reference genome, in this case the GRCh37. Using vcftools[80] (https://vcftools.github.io/index.html), a table containing all of the single nucleotide polymorphisms (SNPs) for each gene was produced and allele frequencies for all sites in each gene were calculated.

Blast (http://blast.ncbi.nlm.nih.gov/Blast.cgi)[81] was used to identify homologous sequences of the human IZUMO1 and JUNO amino acid sequences, totaling 100 different sequences for both proteins. The list of homologous proteins for both IZUMO1 and JUNO were compared to find species that were common to both sets, to investigate whether the two proteins evolved via speciation following the same lines throughout history. In total, a list of 29 species was compiled, and the nucleotide and amino acid sequences for both IZUMO1 and JUNO were retrieved.

### General Statistics

The program SNPeff [26] implemented in Lakehead University’s Galaxy server (https://galaxyproject.org) was used to determine the number of SNPs and their effects. This application determines the number of SNPs, where the SNPs are located and the effects of the SNPs[26, 38]. The SNPeff Genome version name used was GRCh37.74, with an upstream/ downstream length of 5000 bases, with a set size for splice sites in bases of 2 bases and using the default splice Region settings[26]. It should be noted that the IZUMO1 gene is found in the reverse strand.

### Population groups

For population-based analysis, we used the original 1000 Genomes 26 populations, as well as “supergroups” of the original populations (Supplementary Material Table S1). These supergroups were formed along the lines of continental location of the 26 original groups, except that South (SAS) and East (EAS) Asian populations were analyzed individually instead of as a single ASI supergroup. The reason for this, as discussed in the Results section, is that ASI as a supergroup presents variant positions that are not in Hardy-Weinberg equilibrium, whereas as individual groups EAS and SAS have all loci in Hardy-Weinberg equilibrium.

### Population Genetics Parameters and Principal Components Analysis

An R script was developed to calculate population genetics parameters and perform Principal Components Analysis. The script reads vcf files as input and converts the data to the various formats required by individual R packages. The input vcf files are pre-filtered using vcftools[80] to eliminate indels and low frequency SNPs with a MAF specified by the user. MAF filtering, if used, was applied to the entire dataset, as opposed to each or to one of the individual populations. Tajima’s D and nucleotide diversity (π) were calculated using all biallelic sites within the gene segment (i.e., no MAF filtering). Similarly, all variant sites were included in the calculation of µ scores and β statistics. F_ST_ values between each of the regional populations, as well as each of the supergroup populations, were estimated after excluding rare alleles (MAF < 1%), as their inclusion may introduce ascertainment bias if the distribution of rare alleles is uneven across the individual populations[82], which seems to be the case in the 1000 Genomes dataset[83]. LD and haplotype determination were based on variant sites with MAF ≥ 5% to facilitate the identification of haplotypes common to all population groups. The R package vcfR[84] was used to read the vcf files and convert to other formats. Tajima’s D, overall and pairwise Weir and Cockerham[73] F_ST_ values were calculated using the R package HierFSTAT (https://github.com/jgx65/hierfstat). For PCA of F_ST_ values, we used the R package FactoMineR[85]. Geographic population and sex were assigned to each individual sequenced from a table downloaded from http://ftp.1000genomes.ebi.ac.uk/vol1/ftp/technical/working/20130606_sample_info/20130606_sample_info.xlsx. Supergroup membership was assigned according to Supplementary Material Table S1. Principal Components were determined using Weir and Cockerham pairwise F_ST_ values calculated between regional population pairs or their supergroups (these results are presented in the Supplementary Material).

### Haplotype inference

The program Haploview[38] was used to select sites for linkage disequilibrium analysis and haplotype inference. The Galaxy package vcf_to_ped[80] was used to convert the vcf files for IZUMO1 and JUNO into ped files for each of the population group. The vcf files used for haplotype inference for both IZUMO1 and JUNO were filtered to include only SNPs with a MAF of 5% or greater in the entire population set. In Haploview, at the level of population groups, a MAF threshold of 1.0 E-20 was used to select variant sites for further analysis. A minimum number of Mendellian inheritance errors allowed of 1, minimum fraction of nonzero genotypes of 0.75, a Hardy Weinberg p-value threshold of 0.001 and a maximum distance between markers of 10000 in kilobases were also set as input parameters for Haploview.

### Intergenic LD

We used vcftools[80] (https://vcftools.github.io/index.html) to calculate r^2^ between pairs of SNPs where each SNP in the pair came from a different gene. We only considered SNPs with a MAF of 1% or greater. This set, thus, included all SNPs used in Haplotype inference of each gene.

### Structural Analysis of Haplotypes

The PolyPhen-2, a human nonsynonymous SNP prediction database (Polymorphism Phenotyping v2; http://genetics.bwh.harvard.edu/pph2/)[55], was employed to analyzed the resulting amino acid sequences and predict the possible impacts of the amino acid substitutions on the structure and function of IZUMO1 and JUNO. PolyPhen-2 is an automatic tool for predicting the possible impact of amino acid substitutions on the structure and function of a human protein based on the sequence, phylogenetic and structural information characterizing the substitution[55]. The amino acid sequences for each protein were input into the online graphical user interface, the position of the mutation as well as the amino acid substitution were indicated and the software was allowed to analyze the information.

### Ancient Neanderthal Genome Analysis

The ancient genomes of Denisova, Vindija Neanderthal, Altai Neanderthal and Ust’-Ishim ancient modern human were downloaded from the Max Planck Institute for Evolutionary Anthropology Department of Genetics and Bioinformatics Group’s server (http://cdna.eva.mpg.de/neandertal/altai/) in vcf format. The segments corresponding to the IZUMO1 and JUNO genes were isolated using vcftools[86]. Pattern-matching of the DNA markers constituting human haplotypes was used to identify the allele present in these ancient genomes.

### Multiple Sequence Alignment

Pairwise percent identities and multiple sequence alignments were calculated using ClustalX[63]. Amino acid sequences were used to construct an interspecies alignment, whereas an intraspecies alignment was based on whole-gene nucleotide sequences. In the case of human sequences, the percent identity values generated by ClustalX using the vcf files had to be mathematically converted to represent the complete IZUMO1 or JUNO genetic sequences. To do this the number of variant SNPs between the individual and the reference genetic sequence was subtracted from the total number of nucleotides in the gene, that number was then divided by the total number of nucleotides in the gene to get a percent identity for the gene in its entirety.

### Scanning for signals of positive selection

The package RAiSD version 2.9 [64] was used to scan chromosome 11 for signals of positive selection. RAiSD relies on 3 selective sweep signatures to calculate its µ statistic: localized reduction of polymorphisms (µ_var), shift in the Site Frequency Spectrum towards low and high number of derived alleles (µ_SFS), and pattern of linkage disequilibrium (µ_LD) which increases for pairs of loci on the same side of a beneficial mutation but decreases for pairs involving loci on different sides. We choose this application because it implements multiple signatures of selection and it is designed to handle the very large sizes of the datasets corresponding to whole-chromosomes from the 1000 Genomes project. The package was downloaded from https://github.com/alachins/raisd and installed on an in-house Linux computer cluster. The vcf file containing data from the 1000 Genomes project corresponding to human chromosome 11 was downloaded from NCBI (https://www.ncbi.nlm.nih.gov/genome/gdv/). This set contains 3,865,304 SNPs identified from the sequences of 5008 chromatids. Based on recommendation from the authors[64] we omitted from the RAiSD calculations a region corresponding to the centromere in each chromosome. We used a threshold of 0.9995 (99.95%) to identify regions of positive selection as those with µ values above the threshold, whereas values below the threshold are associated to regions of selection neutrality. Genomic position versus beta statistic plots were created using a customized R script.

### Scanning for signals of balancing selection

The package BetaScan2[66] was used to scan chromosome 19 for signals of balancing selection. The package was downloaded from https://github.com/ksiewert/BetaScan and installed on an in-house Linux computer cluster. The vcf files containing data from the 1000 Genomes project corresponding to human chromosome 19 was downloaded from NCBI (https://www.ncbi.nlm.nih.gov/genome/gdv/). This set contains 1,746,975 SNPs. Due to size-handling limitations in the vcf to BetaScan input file converter, glactools[67], we had to split the dataset into population groups, and run the scan for the populations EUR and AFR. We used the top 99.9 percentile as threshold to identify regions of balancing selection as those with β values above the threshold. Genomic position versus beta statistic plots were created using a customized R script.

## Acknowledgements

This work was supported by Natural Sciences and Engineering Research Council of Canada (www.nserc-crsng.gc.ca) grant DDG-2018-00015 to WBF. The funders had no role in study design, data collection and analysis, decision to publish, or preparation of the manuscript. Combined haplotype frequencies were calculated using an R script authored by Dallas Nygard, which is still under development.

## Author contributions

JA performed data collection, analysis, and interpretation, and drafted the manuscript. WBF contributed the original idea and research design, supervised work development and data interpretation, and critically revised the manuscript for submission.

## Supporting Information

Supplementary material is provided and includes:

Table S1 and Table S2: Number of variant sites at each step for IZUMO1 and JUNO, respectively.

Table S3: A list of the 26 different populations sampled by the 1000 Genomes project[23] clustered into five larger population groups.

Table S4: Tajima’s D value analysis of various genes under different types of selection[30–32, 34]. African versus Non-African populations.

Table S5: Hardy-Weinberg Equilibrium analysis of IZUMO1 gene.

Table S6: Hardy-Weinberg Equilibrium analysis of JUNO gene.

Table S7: F_ST_ values in the IZUMO1 gene.

Table S8: F_ST_ values in the JUNO gene.

Table S9: Percent identities in reference to the IZUMO1 *Homo sapiens* amino acid sequence for 29 homologous mammalian species.

Table S10: Percent identities in reference to the JUNO *Homo sapiens* amino acid sequence for 29 homologous mammalian species.

Table S11: The average, maximum and minimum percent identity values for both JUNO and IZUMO1 nucleotide sequences.

Tables S12 and S13: A description of the synonymous and non-synonymous SNPs in the IZUMO1 and the JUNO genes, respectively.

Figures S1 and S2 Histograms of the frequencies of F_ST_ values between all 26 regional populations for the IZUMO1 and the JUNO genes, respectively.

Figure S3 and S4: Principal Component analysis of F_ST_ values between population groups for human IZUMO1 and JUNO genes, respectively.

Figure S5: Scanning for signals of positive selection on chromosome 11 of ALL individuals sequenced in the 1000 Genomes project.

