## Supplementary Material for "Genetic Diversity in the IZUMO1-JUNO Protein-Receptor Pair Involved in Human Reproduction"

*corresponding author

Table S1: Comprehensive breakdown of the variants in the IZUMO1 gene sequence when unfiltered and filtered with a minor allele frequency (MAF) of 5% using SNPEff (1).

| | No maf filtering | Maf 5% frequency |
|---|---|---|
| Variants | 192 | 31 |
| Variant rates | 307963 | 1907386 |
| SNPs | 189 | 31 |
| Insertions | 1 | 0 |
| Deletions | 2 | 0 |
| Low Impact Effects | 199 (29.4%) | 23 (21.5%) |
| Moderate Impact Effects | 11 (1.6%) | 1 (0.9%) |
| Modifier Impact Effects | 466 (68.9%) | 83 (77.6%) |
| Missense Mutations | 12 (57.1%) | 1 (33.3%) |
| Silent Mutations | 9 (42.9%) | 2 (66.7%) |
| Downstream Effects | 143 (21.2%) | 29 (27.1%) |
| Intergenic Effects | 20 (3.0%) | 6 (5.6%) |
| Intragenic Effects | 1 (0.1%) | 0 |
| Intron Effects | 132 (19.5%) | 18 (16.8%) |
| Next Protein Effects | 182 (26.9%) | 21 (19.6%) |
| Non-synonymous Coding Effects | 11 (1.6%) | 1 (0.9%) |
| Non-synonymous Start Effects | 1 (0.1%) | 0 |
| Splice Site Region and Intron Effect | 5 (0.7%) | 0 |
| Start Gained Effect | 2 (0.3%) | 0 |
| Synonymous Coding Effect | 9 (1.3%) | 2 (1.9%) |
| Upstream Effects | 157 (23.3%) | 26 (24.3%) |
| UTR 5 Prime Effect | 13 (1.9%) | 4 (3.7%) |

Table S2: Comprehensive breakdown of the variants in the JUNO gene sequence when unfiltered and filtered with a MAF of 5% using SNPEff (1).

| | No maf filtering | Maf 5% frequency |
|---|---|---|
| Variants | 90 | 7 |
| Variant rates | 1500072 | 19286645 |
| SNPs | 82 | 7 |
| Mixed Variants | 8 | 0 |
| High Impact Effects | 3 (1.8%) | 0 |
| Low Impact Effects | 86 (51.1%) | 3 (30%) |
| Moderate Impact Effects | 19 (11.3%) | 1 (10%) |
| Modifier Impact Effects | 60 (35.7%) | 6 (60%) |
| Missense Mutations | 18 (60%) | 1 (100%) |
| Silent Mutations | 3 (10%) | 0 |
| Downstream Effects | 9 (30%) | 0 |
| Intergenic Effects | 4 (2.4%) | 0 |
| Intron Effects | 53 (31.5%) | 5 (50%) |
| Next Protein Effects | 78 (46.4%) | 3 (30%) |
| Non-synonymous Coding Effects | 18 (10.7%) | 1 (10%) |
| Stop Gained Effects | 3 (1.8%) | 0 |
| Synonymous Coding Effect | 9 (5.4%) | 0 |
| UTR 5 Prime Effect | 3 (1.8%) | 1 (10%) |

Table S3: A list of the 26 different populations sampled by the 1000 Genomes project(2) clustered into five larger population groups, where *n* signifies the number of individuals in each population group.

| **Category** | **n** | **Populations included:** | **Population Code** |
|---|---|---|---|
| South Asia | 489 | Bengali in Bangladesh | BEB |
| | | Gujarati Indian | GIH |
| | | Indian Telugu in the UK | ITU |
| | | Punjabi in Lahore,Pakistan pjl | PJL |
| | | Sri Lankan Tamil in the UK | STU |
| East Asian | 504 | Japanese in Tokyo, Japan | JPT |
| | | Han Chinese in Bejing, | CHB |
| | | China Southern Han Chinese, China | CHS |
| | | Chinese Dai in Xishuangbanna | CDX |
| | | Kinh in Ho Chi Minh City, Vietnam | KHV |
| Europe | 503 | Northern and Western European Finnish in Finland | CEU |
| | | Finnish in Finland | FIN |
| | | British in England and Scotland | GBR |
| | | Iberian populations in Spain | IBS |
| | | Toscani in Italia | TSI |
| America | 347 | Colombian in Medellin, Colombia | CLM |
| | | Mexican Ancestry in Los Angeles, | MXL |
| | | Peruvian in Lima, Peru | PEL |
| | | Puerto Rican in Puerto Rico | PUR |
| Africa | 661 | African Caribbean in Barbados | ACB |
| | | African Ancestry in Southwest US | ASW |
| | | Esan in Nigeria | ESN |
| | | Gambian in Western Division | GWD |
| | | Luhya in Webuye, Kenya | LWK |
| | | Mende in Sierra Leone | MSL |
| | | Yoruba in Ibadan, Nigeria | YRI |

Table S4: Tajima's D analysis of various genes under different types of selection(3-8). Population sizes (n) are reported in parenthesis. Tajima's D was calculated using VCFTools(9) in bins of 100 bp for all biallelic sites within the location range of each gene.

| Selection | Gene | Location (GRCh37.p13) | Tajima's D all populations (n=2,504) | Literature Value | Reference |
|---|---|---|---|---|---|
| Unknown | | | | | |
| | IZUMO1 | Chr 9 49244073-49250831 | -0.35532 | N/A | N/A |
| | JUNO | Chr 11 94038803-94040858 | -0.77916 | N/A | N/A |
| Neutral | | | | | |
| | LTA | Chr 6 31539876-31542101 | -0.45138 | 0.746 (n=282) 6 Chinese populations | (5) |
| | TAS2R38 | Chr 7 141463897-141464997 | -0.58725 | 1.078 (n=8,589) | (10) |
| | TBX1 | Chr 22 19744226-19771116 | -0.69686 | -0.25 (n=124) (22 = EUR, 27 = AFR, 24 = ASI, 22 = AMR) | (11) |
| | VTN | Chr 17 26694298-26697373 | -0.61777 | Value not reported | |
| Balancing | | | | | |
| | ABO | Chr 9 136130563 - 136150630 | -0.07299 | 2.035 EUR (n=23) 1.772 AFR (n=24) | (4, 6) (12) |
| | BPIFB4 | Chr 20 31669318-316699557 | -0.56074 | Value not reported | (7) |
| | BTN1A1 | Chr 6 26500577-26510653 | -0.57767 | Value not reported | (7) |
| | CDSN | Chr 6 31082865- | 0.333637 | Value not reported | (7) |

| | | | | | |
|---|---|---|---|---|---|
| | | 31088252, complement | | | |
| | CLCNKB | Chr 10 16370231-16383821 | -0.34419 | Value not reported | (7) |
| | ERAP2 | Chr 5 96211644-96255420 | -0.30469 | 1.526 (n=180) | (3) |
| | GRIN3A | Chr 9 104331634-104500862, complement | -0.53335 | Value not reported | (7) |
| | HLAA | Chr 6 29910247-29913661 | 0.656452 | 2.9 (n=205) | (7, 13, 14) |
| | HLAB | Chr 6 31321649-31324989, complement | 0.354656 | 2.4 (n=205) | (7, 13, 14) |
| | KRT6C | Chr 12 52862300-52867569, complement | -0.32544 | Value not reported | (7) |
| | KRT84 | Chr 12 52771596-52779417, complement | -0.34531 | Value not reported | (7) |
| | TRIM22 | Chr 11 5710817 - 5732093 | -0.37645 | Value not reported | (7) |
| Positive | | | | | |
| | ABHD1 | Chr 2 27346632-27353680 | -0.72632 | Value not reported | (8) |
| | ALMS1 | Chr 2 73612886-73837047 | -0.67787 | Value not reported | (8) |
| | APOBEC3F | Chr 22 39436609-39451977 | -0.67458 | Value not reported | (8) |
| | APOBEC3G | Chr 22 39473010-39483748 | -0.67032 | Value not reported | (8) |
| | CD36 | Chr 7 80231504-80308593 | -0.5766 | Value not reported | (8) |
| | CD58 | Chr 1 117057156-117113715, complement | -0.70801 | Value not reported | (8) |

| | | | | | |
|---|---|---|---|---|---|
| | CD72 | Chr 9 35609976 -35618862, complement | -0.7711 | Value not reported | (8) |
| | EDAR | Chr 2 109510927-109605828 | -0.66222 | Value not reported | (8, 15) |
| | FAF1 | Chr 1 50906935-51425936, complement | -0.73988 | Value not reported | (8) |
| | GRAP2 | Chr 22 40297086 - 40369347 | -0.7289 | Value not reported | (8) |
| | HYAL3 | Chr 3 50330259 - 50336899, complement | -0.73139 | Value not reported | (8) |
| | ITGAE | Chr 17 3617919 - 3704537, complement | -0.57616 | Value not reported | (8) |
| | KEL | Chr 7 142638201-142659503 | -0.76554 | -2.467 EUR (n=23) -0.823 AFR (n=24) | (6) |
| | LCT | Chr 2 136545415-136594750, complement | -0.61215 | Value not reported | (16) |
| | NPAP1 | Chr 15 24920541-24928593 | -0.74745 | Value not reported | (8) |
| | PRM1 | Chr 16 11374693-11375192, complement | -0.84476 | Value not reported | (8) |
| | PRM2 | Chr 16 11369493-11370337, complement | -0.91328 | Value not reported | (8) |
| | PROS1 | Ch3 3 93591881-93692934, complement | -0.80824 | -1.44 (n = 47) (24 = AFR, 23 = EUR) | (17) |

| | | | | | |
|---|---|---|---|---|---|
| | RBAK | Chr 7 5085452-5112854 | -0.62683 | Value not reported | (8) |
| | RHCE | Chr 1 25687853-25747363, complement | -0.63686 | Value not reported | (8) |
| | SLC24A5 | Chr 15 48413169-48434926 | -0.73307 | Value not reported | (8) |
| | SYT1 | Chr 12 79257773-79845788 | -0.66917 | Value not reported | (8) |
| | TRPV6 | Chr 7 142568956-142583490 | -0.67565 | -2.865 EUR (n=23) 0.893 AFR (n=24) | (6, 18, 19) |

Table S5: Hardy-Weinberg Equilibrium analysis of IZUMO1 gene in males only and the entire population in each of the groups included in the analyzed haplotype(20).

| Location | rs2307018 | | rs2307019 | | rs838148 | |
|---|---|---|---|---|---|---|
| Population | All | Males | All | Males | All | Males |
| AFR | 0.7809 | 1 | 0.7809 | 1 | 0.8287 | 0.9697 |
| AMR | 0.9762 | 0.57 | 0.9762 | 0.57 | **0.9762** | **0.2741** |
| EUR | 0.3691 | 1 | 0.3691 | 1 | 0.8109 | 0.428 |
| EAS | 0.8668 | 0.7768 | 0.8668 | 0.7768 | 0.9064 | 1.00 |
| SAS | 0.3308 | 0.2600 | 0.3308 | 0.2600 | 1 | 0.8749 |
| ASI | **5.83E-06** | **6.91E-05** | **5.83E-06** | **6.91E-05** | 0.3669 | 0.3098 |
| ALL | 2.06E-13 | 4.62E-08 | 2.06E-13 | 4.62E-08 | 0.0039 | 0.0025 |

Table S6: Hardy-Weinberg Equilibrium analysis of JUNO gene in females only and the entire population in each of the groups included in the analyzed haplotype(20).

| Location | rs61742524 | | rs55784852 | | rs16920146 | | rs7925833 | | rs7935583 | |
|---|---|---|---|---|---|---|---|---|---|---|
| Population | All | Females | All | Females | All | Females | All | Females | All | Females |
| AFR | **0.4875** | **0.0221** | **0.4875** | **0.022** | **0.4875** | **0.0221** | **0.666** | **0.1764** | **0.4875** | **0.0221** |
| AMR | **0.491** | **0.1201** | **0.491** | **0.120** | **0.491** | **0.1201** | **0.4049** | **0.0725** | **0.491** | **0.1201** |
| EUR | 1 | 1 | 1 | 1 | 1 | 1 | 1 | 1 | 1 | 1 |
| EAS | 1 | 1 | 1 | 1 | 1 | 1 | 1 | 1 | 1 | 1 |
| SAS | 1 | 1 | 1 | 1 | 1 | 1 | 1 | 1 | 1 | 1 |
| ASI | 1 | 1 | 1 | 1 | 1 | 1 | 1 | 1 | 1 | 1 |
| ALL | 1.36E-20 | 4.25E-10 | 5.26E-20 | 4.25E-10 | 5.26E-20 | 4.25E-10 | 1.17E-20 | 009.28E-06 | 1.78E-20 | 4.25E-10 |

Table S7: FST values in the IZUMO1 gene between the five larger population groups for the entire set of 2504 individuals sampled in the 1000 Genomes project. These FST values were calculated using SNPs with a MAF of at least 1%. For comparison, a genome wide FST value for the human genome is 0.12. The average of all pairwise values is 0.150.

| | EUR | EAS | AMR | SAS | AFR |
|---|---|---|---|---|---|
| EUR | | 0.296 | 0.023 | 0.020 | 0.080 |
| EAS | 0.296 | | 0.196 | 0.224 | 0.447 |
| AMR | 0.023 | 0.196 | | 0.004 | 0.123 |
| SAS | 0.020 | 0.224 | 0.004 | | 0.085 |
| AFR | 0.080 | 0.447 | 0.123 | 0.085 | |

Table S8: FST values in the JUNO gene between the five larger population groups for the entire set of 2504 individuals sampled in the 1000 Genomes project. These FST values were calculated using SNPs with a MAF of at least 1%. For comparison, a genome wide FST value for the human genome is 0.12. The average of all pairwise values is 0.135.

| | EUR | EAS | AMR | SAS | AFR |
|---|---|---|---|---|---|
| EUR | | 0.001 | 0.007 | 0.114 | 0.310 |
| EAS | 0.001 | | 0.004 | 0.096 | 0.304 |
| AMR | 0.007 | 0.004 | | 0.063 | 0.247 |
| SAS | 0.114 | 0.096 | 0.063 | | 0.203 |
| AFR | 0.310 | 0.304 | 0.247 | 0.203 | |

Table S9: Percent identities in reference to the IZUMO1 *Homo sapiens* amino acid sequence for 29 homologous mammalian species. The E-value indicates the statistical significance of the data, the smaller the number the better, and the query cover indicates the percentage of the sequence that overlaps with the *Homo sapiens* sequence. The % identity BLAST is generated by a local alignment and were acquired from http://www.ncbi.nlm.nih.gov.The % identity ClustalX is generated by global alignment and was calculated using ClustalX(21).

| **Species** | **Common Name** | **E-value** | **Query Cover** | **% Identity BLAST** | **% Identity Clustalx** |
|---|---|---|---|---|---|
| Homo sapiens | Human | 2.00E-164 | 100 | 100 | 100 |
| Pan troglodytes | Chimpanzee | 4.00E-92 | 100 | 100 | 99 |
| Gorilla gorilla gorilla | Lowland Gorilla | 3.00E-161 | 100 | 99 | 98 |
| Nomascus leucogenys | White-cheeked gibbon | 3.00E-155 | 100 | 95 | 94 |
| Papio Anubis | Baboon | 6.00E-148 | 100 | 92 | 91 |
| Mandrillus leucophaeus | Drill | 2.00E-147 | 100 | 92 | 91 |
| Macaca fascicularis | Long-tailed macaque | 3.00E-148 | 100 | 92 | 89 |
| Macaca nemestrina | Pigtail monkey | 4.00E-150 | 100 | 92 | 92 |
| Rhinopithecus roxellana | Golden snub-nosed monkey | 3.00E-146 | 100 | 91 | 91 |
| Colobus angolensis palliates | Peter’s Angola Colobus | 3.00E-147 | 100 | 92 | 91 |
| Saimiri boliviensis boliviensis | Black-headed squirrel monkey | 3.00E-125 | 94 | 83 | 81 |
| Aotus nancymaae | Nancy Ma’s night monkey | 4.00E-129 | 100 | 82 | 84 |
| Propithecus coquereli | Coqueral’s sifaka | 7.00E-108 | 98 | 71 | 70 |
| Felis catus | Cat | 9.00E-76 | 96 | 58 | 50 |
| Microcebus murinus | Gray mouse lemur | 8.00E-87 | 98 | 59 | 64 |
| Equus przewalskii | Przewalski horse | 1.00E-89 | 97 | 66 | 59 |
| Pteropus vampyrus | Large flying fox (bat) | 8.00E-80 | 100 | 56 | 58 |
| Otolemur garnettii | Northern Greater Galago | 4.00E-81 | 82 | 56 | 56 |
| Loxodonta Africana | African savannah elephant | 2.00E-84 | 74 | 72 | 56 |
| Camelus dromedaries | One-humped camel | 2.00E-94 | 100 | 63 | 65 |
| Trichechus manatus latirostris | West indian manatee | 2.00E-84 | 67 | 76 | 55 |
| Orcinus orca | Orca | 1.00E-96 | 99 | 64 | 65 |
| Heterocephalus glaber | Naked mole rat | 5.00E-85 | 100 | 60 | 55 |
| Microtus ochrogaster | Prairie voles | 5.00E-80 | 75 | 63 | 52 |
| Chinchilla lanigera | Long-tailed chinchilla | 1.00E-85 | 100 | 64 | 59 |
| Jaculus jaculus | Lesser Egyptian Jerboa | 1.00E-79 | 71 | 67 | 53 |
| Rattus norvegicus | Rat | 1.00E-79 | 75 | 63 | 55 |
| Octodon degus | Degu | 1.00E-82 | 73 | 69 | 55 |
| Bos mutus | Wild yak | 1.00E-84 | 89 | 64 | 60 |
| Mus musculus | Mouse | 1.00 E-81 | 78 | 46 | 47 |

Table S10: Percent identities in reference to the JUNO *Homo sapiens* amino acid sequence for 29 homologous mammalian species. The E-value indicates the statistical significance of the data, the smaller the number the better, and the query cover indicates the percentage of the sequence that overlaps with the *Homo sapiens* sequence. The % identity BLAST is generated by a local alignment and were acquired from http://www.ncbi.nlm.nih.gov.The % identity ClustalX is generated by global alignment and was calculated using ClustalX(21).

| **Species** | **Common Name** | **E-value** | **Query Cover** | **% Identity BLAST** | **% Identity Clustalx** |
|---|---|---|---|---|---|
| Homo sapiens | Human | 7.00E-173 | 100 | 100 | 100 |
| Pan troglodytes | Chimpanzee | 1.00E-163 | 95 | 100 | 99 |
| Gorilla gorilla gorilla | Lowland Gorilla | 2.00E-163 | 95 | 99 | 99 |
| Nomascus leucogenys | White-cheeked gibbon | 1.00E-160 | 95 | 98 | 97 |
| Papio Anubis | Baboon | 2.00E-159 | 95 | 97 | 96 |
| Mandrillus leucophaeus | Drill | 5.00E-159 | 95 | 97 | 96 |
| Macaca fascicularis | Long-tailed macaque | 5.00E-158 | 95 | 96 | 96 |
| Macaca nemestrina | Pigtail monkey | 3.00E-158 | 95 | 96 | 95 |
| Rhinopithecus roxellana | Golden snub-nosed monkey | 6.00E-156 | 93 | 97 | 95 |
| Colobus angolensis palliates | Peter’s Angola Colobus | 1.00E-152 | 93 | 96 | 94 |
| Saimiri boliviensis boliviensis | Black-headed squirrel monkey | 6.00E-150 | 95 | 92 | 92 |
| Aotus nancymaae | Nancy Ma’s night monkey | 9.00E-148 | 95 | 90 | 90 |
| Propithecus coquereli | Coqueral’s sifaka | 8.00E-145 | 100 | 87 | 86 |
| Felis catus | Cat | 4.00E-132 | 99 | 77 | 77 |
| Microcebus murinus | Gray mouse lemur | 2.00E-131 | 100 | 80 | 82 |
| Equus przewalskii | Przewalski horse | 5.00E-122 | 95 | 77 | 77 |
| Pteropus vampyrus | Large flying fox (bat) | 3.00E-121 | 94 | 78 | 79 |
| Otolemur garnettii | Northern Greater Galago | 2.00E-119 | 94 | 76 | 75 |
| Loxodonta Africana | African savannah elephant | 1.00E-116 | 94 | 76 | 76 |
| Camelus dromedaries | One-humped camel | 7.00E-116 | 93 | 78 | 76 |
| Trichechus manatus latirostris | West indian manatee | 4.00E-116 | 92 | 76 | 77 |
| Orcinus orca | Orca | 5.00E-115 | 94 | 76 | 74 |
| Heterocephalus glaber | Naked mole rat | 5.00E-115 | 99 | 71 | 73 |
| Microtus ochrogaster | Prairie voles | 4.00E-112 | 98 | 73 | 74 |
| Chinchilla lanigera | Long-tailed chinchilla | 2.00E-112 | 100 | 71 | 72 |
| Jaculus jaculus | Lesser Egyptian Jerboa | 8.00E-109 | 99 | 72 | 71 |
| Rattus norvegicus | Rat | 1.00E-105 | 98 | 69 | 69 |
| Octodon degus | Degu | 1.00E-101 | 98 | 66 | 67 |
| Bos mutus | Wild yak | 6.00E-101 | 88 | 72 | 68 |
| Mus musculus | Mouse | 8.00E-103 | 99 | 67 | 66 |

Table S11: The average, maximum and minimum percent identity values for both JUNO and IZUMO1 nucleotide sequences for all of the 2504 individuals in reference to the GRCh37 reference genome used in the 1000 Genomes project.

| **Protein** | **JUNO** | **IZUMO1** |
|---|---|---|
| Average %ID | 99.89 | 99.50 |
| Max %ID | 100.00 | 100.00 |
| Min %ID | 99.49 | 99.25 |
| Standard Deviation | 0.11 | 0.24 |

Table S12: A description of the synonymous and non-synonymous SNPs in the IZUMO1 gene when filtered by a MAF of 5%.

| SNP | rs2307018 | rs2307019 | rs8108468 |
|---|---|---|---|
| Effect | Synonymous Coding | Non-synonymous Coding | Synonymous Coding |
| | Next Protein Effect | Next Protein Effect | Next Protein Effect |
| Impact | Low Impact | Moderate Impact | Low Impact |
| Location | Topological Domain: Cytoplasmic | Topological Domain: Cytoplasmic | Topological Domain: Extracellular |
| Amino Acid | A333 | A33V | F107 |
| | Upstream Modifier (RASIP1) | Upstream Modifier (RASIP1) | Upstream Modifier (RASIP1) Downstream Modifier (FUT1) |
| Type of Mutation | Silent Mutation | Missense Mutation | Silent Mutation |

Table S13: A description of the synonymous and non-synonymous SNPs in the JUNO gene when filtered by a MAF of 5%.

| SNP | rs61742524 |
|---|---|
| Effect | Non-synonymous Coding |
| | Next Protein Effect |
| Impact | Moderate Impact |
| Location | Topological Domain: Cytoplasmic |
| Amino Acid | C3W |
| Type of Mutation | Missense Mutation |

Figure S1: Histograms of the frequencies of FST values between all 26 regional populations for the IZUMO1 gene between A) all 2504 individuals sampled in the 1000 Genomes Project, B) just the males sampled and C) just the females sampled. The red line indicates the 0.102 reference value for average human genome-wide FST(22). The maximum FST value was 0.503 between the YRI and CDX populations. For just the male population the maximum FST value was 0.654 between the YRI and CDX populations. For just the female population the maximum FST value was 0.528 between the MSL and CDX populations. These values were calculated using all SNPs with a MAF of 1% or greater.

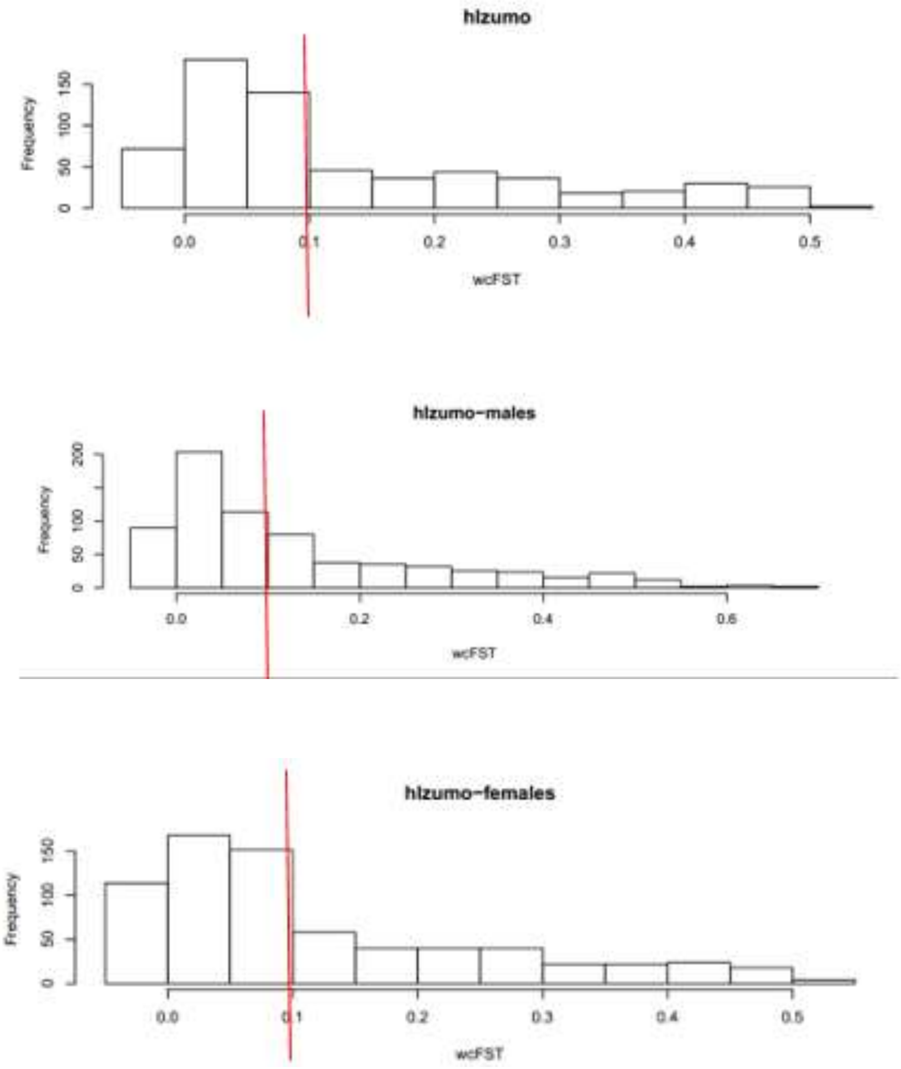

Figure S2: Histograms of the frequencies of FST values between all 26 regional populations for the JUNO gene between A) all 2504 individuals sampled in the 1000 Genomes Project, B) just the males sampled and C) just the females sampled. The red line indicates the 0.102 reference value for average human genome-wide FST(22). The maximum FST value was 0.372 between the MSL and CDX populations. For just the male population the maximum FST value was 0.393 also between the MSL and CDX populations. For just the female population the maximum FST value was 0.528 between the MSL and CDX populations. These values were calculated using all of the SNPs with a MAF of 1% or greater.

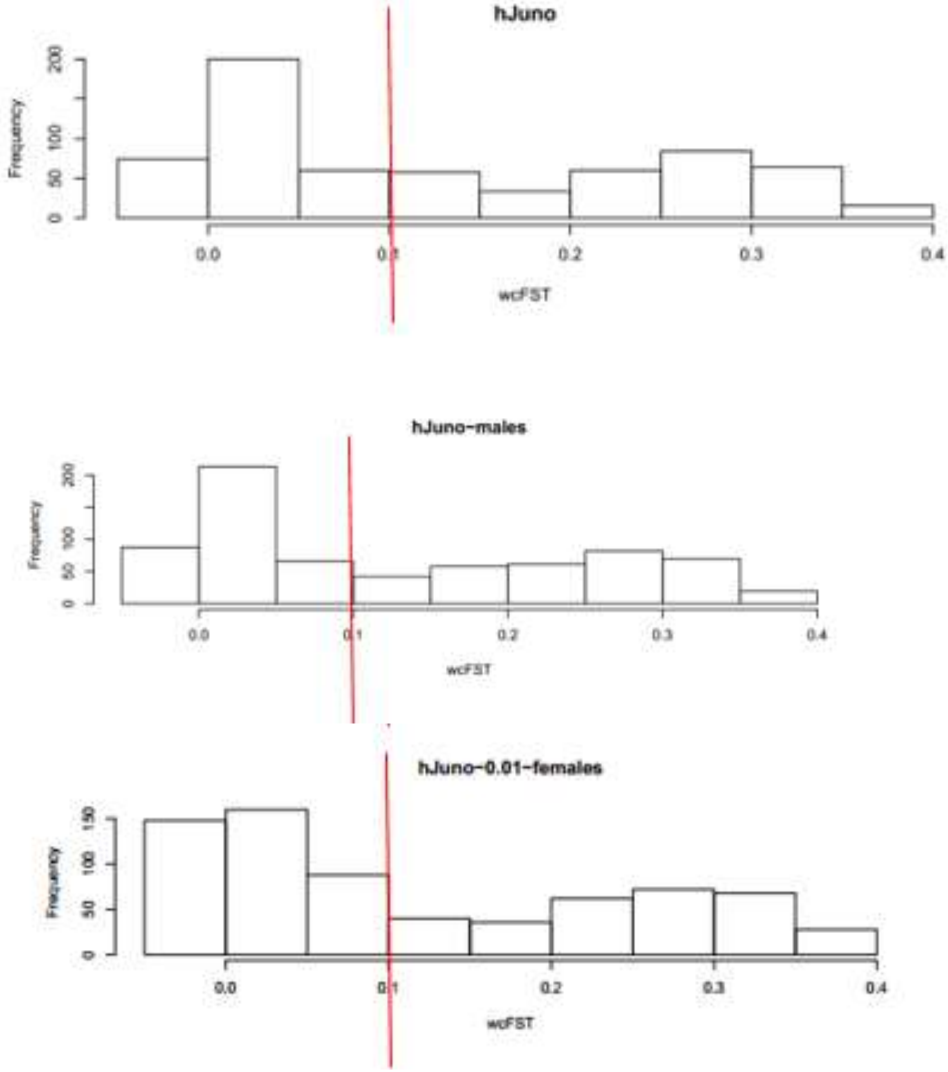

Figure S3: Principal Component analysis of FST values between population groups for human IZUMO1 calculated for the entire set of 2,504 individuals sampled in the 1000 Genomes project. Squares are populations categorized in the supergroup AFR (African); crosses are EUR (European); circles are EAS (East Asian); stars are SAS (South Asian); triangles are AMR (American). The population designations follow the 1000 Genome project annotations, as indicated in Table S3. For human Izumo1, the supergroups AFR and EAS segregate from the others. Components 1 and 2 carry 99% of the information contained in the pairwise FST values and are, thus, effective proxies of the genetic diversity between the populations studied.

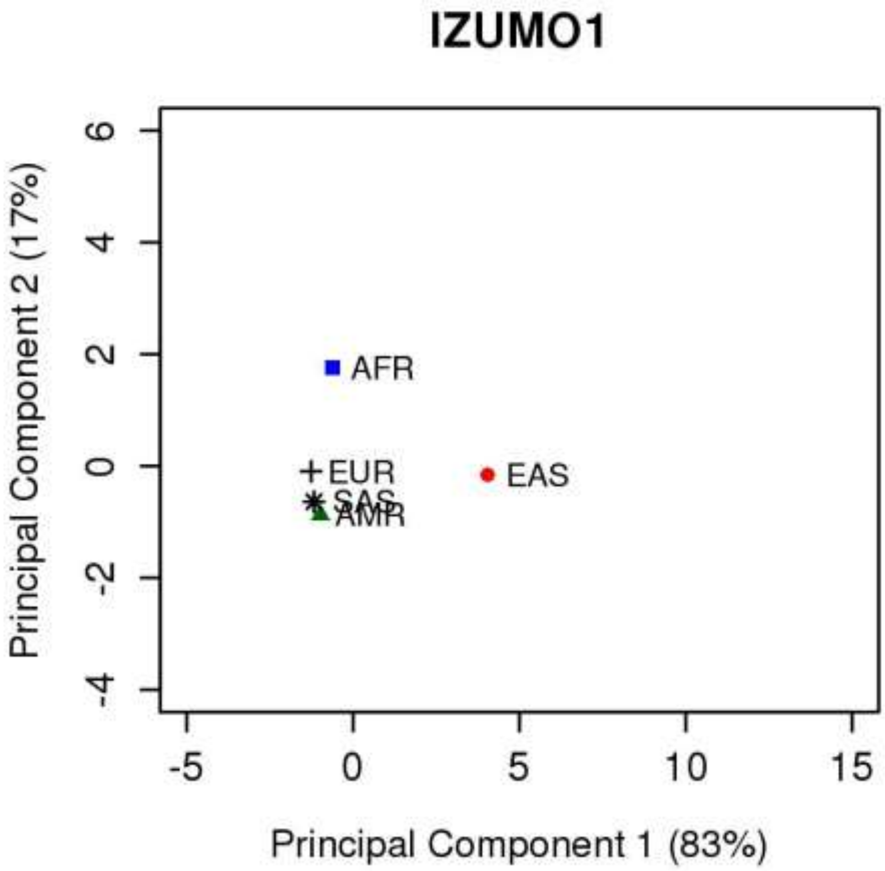

Figure S4: Principal Component analysis of FST values between population groups for human JUNO calculated for the entire set of 2504 individuals sampled in the 1000 Genomes project. Squares are populations categorized in the supergroup AFR (African); crosses are EUR (European); circles are EAS (East Asian); stars are SAS (South Asian); triangles are AMR (American). The population designations follow the 1000 Genome project annotations, as indicated in Table S3. For human Juno, the supergroups AFR and SAS segregate from the others. Components 1 and 2 carry 100% of the information contained in the pairwise FST values and are, thus, perfect proxies of the genetic diversity between the populations studied.

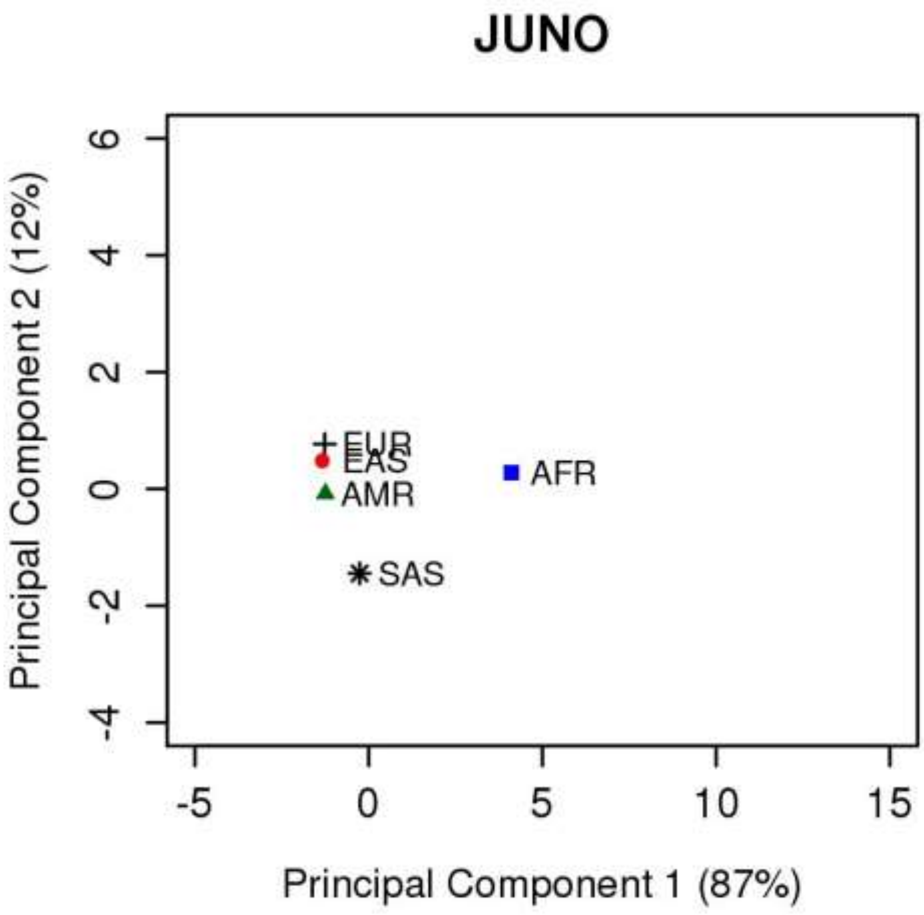

Figure S5: Scanning for signals of positive selection on chromosome 11 of ALL individuals sequenced in the 1000 Genomes project. Regions with μ scores above the 99.95% threshold (solid red line) are expected to be under positive selection. This threshold is based on all μ scores for a dataset and it is, hence, population-dependent. The genomic region corresponding to the JUNO gene is marked by green dashed lines. the genomic region corresponding to JUNO is within a region of μ values modestly above the threshold set and borders a region of high μ values. This suggests that the indicators of positive selection we identified in JUNO from the analysis of nucleotide diversity, Tajima's D, and haplotype inference could have originated from a hitchhiking effect of a selective sweep (23).

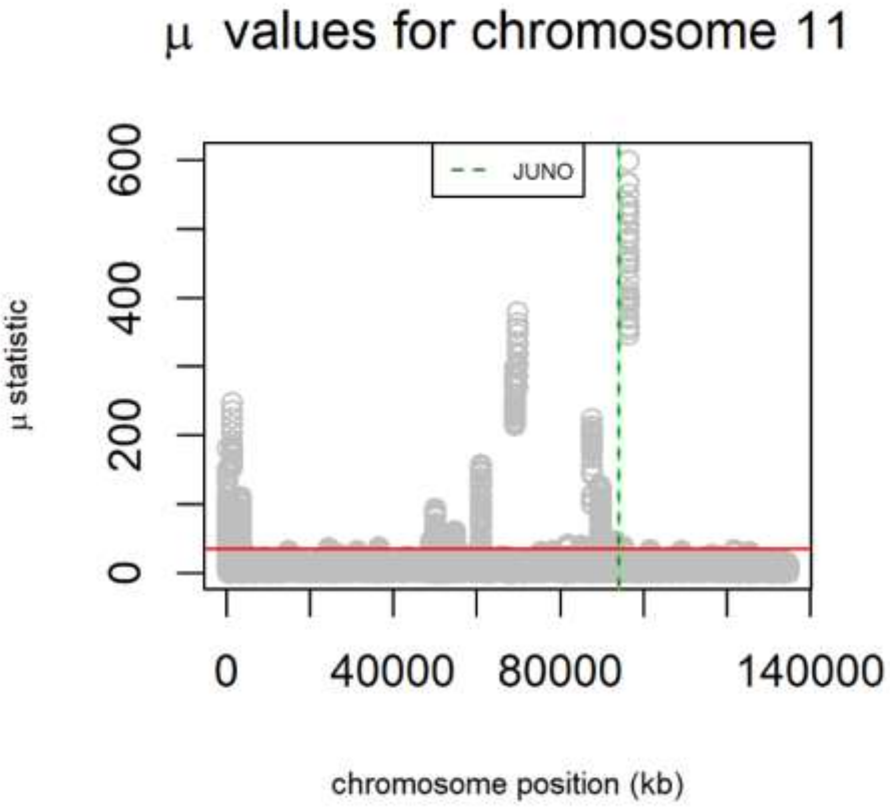

TBX1 in Individuals with and without 22q11.2

Deletion Syndrome. Birth Defects Research (Part A). 2010;88(1):54-63.
12. Saitou N YF. Evolution of primate ABO blood group genes and their homologous genes. Mol Biol Evol. 1997;14(4):399-411.
13. Goeury T, Creary LE, Brunet L, Galan M, Pasquier M, Kervaire B, et al. Deciphering the fine nucleotide diversity of full HLA class I and class II genes in a well-documented population from sub-Saharan Africa. HLA. 2018;91(1):36-51.
14. Hedrick PW, Thomson G. Evidence for Balancing Selection at Hla. Genetics. 1983;104(3):449-56.
15. Bryk Jea. Positive selection in East Asians for an EDAR allele that enhances NF-kappaB activation. . PLos One. 2008;3(5).
16. Bersaglieri T, Sabeti PC, Patterson N, Vanderploeg T, Schaffner SF, Drake JA, et al. Genetic signatures of strong recent positive selection at the lactase gene. American Journal of Human Genetics. 2004;74(6):1111-20.
17. Reed FA, Akey JM, Aquadro CF. Fitting background-selection predictions to levels of nucleotide variation and divergence along the human autosomes. Genome Res. 2005;15(9):1211-21.
18. Korneliussen TS, Moltke I, Albrechtsen A, Nielsen R. Calculation of Tajima's D and other neutrality test statistics from low depth next-generation sequencing data. BMC Bioinformatics. 2013;14(289):289.
19. Hughes DA, Tang K, Strotmann R, Schoneberg T, Prenen J, Nilius B, et al. Parallel selection on TRPV6 in human populations. PLoS One. 2008;3(2):e1686.

20. Barrett JC, Fry B, Maller J, Daly MJ. Haploview: analysis and visualization of LD and haplotype maps. Bioinformatics. 2005;21(2):263-5.
21. Larkin MA, Blackshields G, Brown NP, Chenna R, McGettigan PA, McWilliam H, et al. Clustal W and Clustal X version 2.0. Bioinformatics. 2007;23(21):2947-8.
22. Bhatia G, N. Patterson, et al. . Estimating and interpreting F-ST: The impact of rare variants. Genome Res. 2013;23(9):1514-21.
23. Smith JM, Haigh J. The hitch-hiking effect of a favourable gene. Genet Res. 1974;23(1):23-35.
